# End-of-life targeted auxin-mediated degradation of DAF-2 Insulin/IGF-1 receptor promotes longevity free from growth-related pathologies

**DOI:** 10.1101/2021.05.31.446422

**Authors:** Richard Venz, Tina Pekec, Iskra Katic, Rafal Ciosk, Collin Y. Ewald

**Affiliations:** Eidgenössische Technische Hochschule Zürich, Department of Health Sciences and Technology, Institute of Translational Medicine, Schwerzenbach-Zürich CH-8603, Switzerland; University of Basel, Faculty of Natural Sciences, Klingelbergstrasse 70, 3026 Basel, Switzerland; Friedrich Miescher Institute for Biomedical Research, Maulbeerstrasse 66, 4058 Basel, Switzerland; Institute of Bioorganic Chemistry, Polish Academy of Sciences, Noskowskiego 12/14, 61-704 Poznań, Poland; University of Oslo, Department of Biosciences, Blindernveien 31, 0371 Oslo, Norway

**Keywords:** Insulin, IGF-1, *daf-2*, *daf-16*, *skn-1*, starvation, diet, auxin-induced degradation, neuron, aging, stress, healthspan, dauer

## Abstract

Preferably, lifespan-extending therapies should work when applied late in life without causing undesired pathologies. However, identifying lifespan-extending interventions that are effective late in life and which avoid undesired secondary pathologies remains elusive. Reducing Insulin/IGF-1 signaling (IIS) increases lifespan across species, but the effects of reduced IIS interventions in extreme geriatric ages remains unknown. Using the nematode *C. elegans*, we engineered the conditional depletion of the DAF-2/insulin/IGF-1 transmembrane receptor using an auxin-inducible degradation (AID) system that allows for the temporal and spatial reduction in DAF-2 protein levels at time points after which interventions such as RNAi may lose efficacy. Using this system, we found that AID-mediated depletion of DAF-2 protein efficiently extends animal lifespan. Depletion of DAF-2 during early adulthood resulted in multiple adverse phenotypes, including growth retardation, germline shrinkage, egg-retention, and reducing offspring. By contrast, however, AID-mediated depletion of DAF-2 specifically in the intestine resulted in an extension of lifespan without these deleterious effects. Importantly, AID-mediated depletion of DAF-2 protein in animals past their median lifespan allowed for an extension of lifespan without affecting growth or behavioral capacity. Thus, both late-in-life targeting and tissue-specific targeting of IIS minimize the deleterious effects typically seen with interventions that reduced IIS, suggesting potential therapeutic methods by which longevity and healthspan can be increased in even geriatric populations.

## Introduction

The goal of aging research or geroscience is to identify interventions that promote health during old age (B. K. Kennedy et al., 2014; López-Otín et al., 2013; Partridge et al., 2018). Nutrient-sensing pathways that regulate growth and stress resistance play major roles as conserved assurance pathways for healthy aging (Kenyon, 2010; Liu et al., 2020; López-Otín et al., 2013; Partridge et al., 2018). One of the first longevity pathways discovered was the insulin/insulin-like growth factor (IGF)-1 signaling pathway (reviewed in (Kenyon, 2010)). Reducing insulin/IGF-1 signaling (IIS) increases lifespan across species (Kenyon, 2010). Mice heterozygous for the IGF-1 receptor, or with depleted insulin receptor in adipose tissue, are stress-resistant and long-lived (Blüher et al., 2003; Holzenberger et al., 2003), for example, and several single-nucleotide polymorphisms in the IIS pathway have been associated with human longevity (Kenyon, 2010). Moreover, gene variants in the IGF-1 receptor have been associated and functionally linked with long lifespans in human centenarians (Suh et al., 2008). This suggests that a comprehensive understanding of this pathway in experimental, genetically-tractable organisms has promising translational value for promoting health in elderly humans. However, whether or not reducing insulin/IGF-1 signaling during end-of-life stages can still promote health and longevity in any organism is unknown. Therefore, we turned to the model organism *Caenorhabditis elegans* to investigate whether reducing IIS during old age was sufficient to increase lifespan.

The groundbreaking discovery that a single mutation in *daf-2*, which is the orthologue of both the insulin and IGF-1 receptors (Kimura et al., 1997), or mutations in “downstream” genes in the insulin/IGF-1 signaling pathway, could double the lifespan of an organism was made in the nematode *C. elegans* (Friedman and Johnson, 1988; Kenyon et al., 1993). Since its discovery, over 1000 papers on *daf-2* have been published, making it one of the most studied genes in this model organism (Source: PubMed). Genetic and-omics approaches have revealed that the DAF-2 insulin/IGF-1 receptor signaling regulates growth, development, metabolism, inter-tissue signaling, immunity, stress defense, and protein homeostasis, including extracellular matrix remodeling (Ewald et al., 2015; Gems et al., 1998; Kimura et al., 1997; Murphy, 2013; Wolkow et al., 2000). Much of our knowledge of the effects of *daf-2* on aging has come from the study of reduction-of-function alleles of *daf-2*. Several alleles of *daf-2* have been isolated that are temperature-sensitive with respect to an alternative developmental trajectory. For instance, most *daf-2* mutants develop into adults at 15°C and 20°C but enter the dauer stage at 25°C (Gems et al., 1998), a facultative and alternative larval endurance stage in which *C. elegans* spends most of its life cycle in the wild (Hu, 2007). Under favorable conditions, *C. elegans* develops through four larval stages (L1-L4). By contrast, when the animals are deprived of food, experience thermal stress (above 27°C), or overcrowded environments, the developing larvae molt into an alternative pre-dauer (L2d) stage. If conditions do not improve, *C. elegans* enter the dauer diapause instead of the L3 stage (Hu, 2007; Karp, 2018).

A major limitation in using *daf-2* mutants is that all of them go through this alternative pre-dauer stage (L2d), irrespective of temperature or whether they later enter the dauer stage or not (Karp, 2018; Ruaud et al., 2011). This suggests a physiological reprogramming occurs in these mutants that might spill over to affect physiology in adulthood. Indeed, the *daf-2* alleles have been categorized into two mutant classes depending on the penetrance of dauer-associated phenotypes during adulthood and aging, such as reduced brood size, small body size, and germline shrinkage ^12,15,19-23^. To overcome this, RNA interference of *daf-2* can be applied, which increases lifespan without dauer formation during development and circumvents induction of dauer-associated phenotypes during adulthood (Dillin et al., 2002; Ewald et al., 2015; 2018; S. Kennedy et al., 2004). However, the increase in lifespan by RNAi of *daf-2* is only partial compared to strong alleles such as *daf-2(e1370)* (Ewald et al., 2015). Furthermore, adult-specific RNAi knockdown of *daf-2* quickly loses its potential to increase lifespan and does not extend lifespan when started after day 6 of adulthood (Dillin et al., 2002), after the reproductive period of *C. elegans*. Whether this is due to age-related functional decline of RNAi machinery or residual DAF-2 protein levels -- or whether the late-life depletion of *daf-2* simply does not extend lifespan -- remains unclear. As such, use of an alternative method to reduce DAF-2 levels, beyond RNAi or *daf-2* mutation may allow us to more clearly uncouple the pleiotropic effects of reduced IIS during development from those that drive *daf-2*-mediated longevity during late adulthood.

To this end, we used an auxin-inducible degradation (AID) system to induce the depletion of the degron-tagged DAF-2 protein with temporal precision (Zhang et al., 2015). The *Arabidopsis thaliana* IAA17 degron is a 68-amino acid motif that is specifically recognized by the transport inhibitor response 1 (TIR1) protein only in the presence of the plant hormone auxin (indole-3-acetic acid) (Dharmasiri et al., 2005). Although cytoplasmic, nuclear, and membrane-binding domain proteins tagged with degron have been recently shown to be targeted and degraded in *C. elegans* (Beer et al., 2019; Zhang et al., 2015), to our knowledge, the AID system has not been used previously to degrade transmembrane proteins, such as the DAF-2 insulin/IGF-1 receptor. We find that using AID effectively degrades DAF-2 protein and promotes dauer formation when applied early in development. Dauer-associated phenotypes are present in adults when AID of DAF-2 is applied late in development. Some of these adulthood dauer-traits are induced by the loss of *daf-2* in neurons, but others seem to be caused by the systemic loss of *daf-2*. More importantly, the post-developmental, conditional degradation of DAF-2 protein extends lifespan without introducing dauer-like phenotypes. Remarkably, we demonstrate that when half of the population has died at day 25 of adulthood, AID of DAF-2 in these remaining and aged animals is sufficient to promote longevity. Our work suggests that therapeutics applied at even extremely late stages of life are capable of increasing longevity and healthspan in animals.

## Results

### Generation and validation of a degron-tagged DAF-2 receptor

To monitor and conditionally regulate protein levels of the *C. elegans* DAF-2 insulin/IGF-1 receptor, we introduced a degron::3xFLAG tag into the 3’ end of the *daf-2* open reading frame (Supplementary Figure 1A). This degron::3xFLAG insertion into the genome was designed to tag the DAF-2 receptor at the cytosolic part for two reasons: first, to minimize any interference by the 81-amino acids large degron::3xFLAG-tag with the DAF-2 receptor function; and second, to ensure accessibility of the degron for targeted degradation by the TIR1 ubiquitin ligase expressed in the cytoplasm (Figure 1A). Using CRISPR, we endogenously tagged the DAF-2 receptor and the resulting *daf-2(bch40)* CRISPR allele was verified by PCR (Supplementary Figure 1B). We performed western blot analysis against the 3xFLAG-tag and detected a specific band in *daf-2(bch40)* animals. This band was absent in wild type (N2) and animals carrying only the *eft-3p*::TIR1::mRuby::*unc-54* 3’UTR transgene, which expresses TIR1 in all somatic cells (Figure 1B). To promote degradation of the degron::3xFLAG-tagged DAF-2 receptor, we crossed *daf-2(bch40)* into TIR1-expressing *C. elegans* (Figure 1A). The strain obtained from this cross will be called„ DAF-2::degron” throughout this paper (*ieSi57* [P*eft-3*::TIR1::mRuby::*unc-54* 3’UTR + *Cbr-unc-119*(+)] II; *daf-2(bch40* [degron::3xFLAG::STOP::SL2-SV40-degron::wrmScarlet-*egl-13* NLS]) III.). This strain showed no obvious phenotypes and exhibited a normal developmental progression at 20°C (Supplementary Figures 1C, 1D). To verify whether the band from the western blot was indeed DAF-2::degron::3xFLAG, we treated DAF-2::degron animals with *daf-2* RNAi. The band nearly completely disappeared after 48 hours of *daf-2(RNAi)* feeding (Figure 1C, Data Source Files 1, 2). Collectively, these results suggested that the tagged transmembrane receptor DAF-2 did not interfere with normal DAF-2 function.

**Figure 1.**
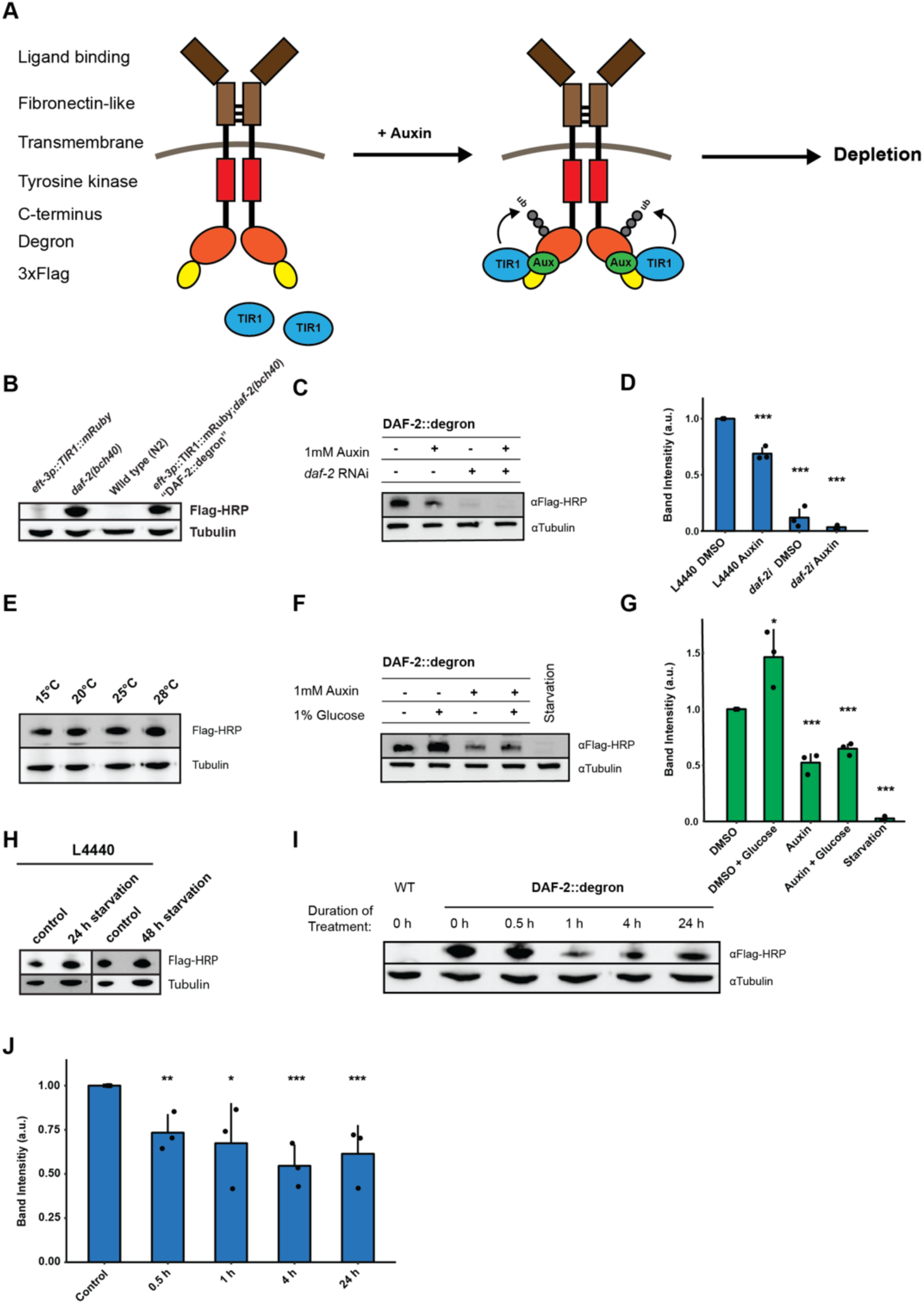
Degron-tagged DAF-2 is functional and susceptible to auxin-mediated degradation. (A) Schematic illustration of AID-mediated DAF-2 receptor depletion in *daf-2(bch40) C. elegans*. (B) Immunoblot analysis of *eft-3p::TIR1::mRuby::unc-54* 3’UTR, *daf-2(bch40)*, wild type (N2) and DAF-2::degron (*eft-3p::TIR1::mRuby::unc-54* 3’UTR; *daf-2(bch40)*). (C) Immunoblot analysis of “DAF-2::degron*”* animals after treatment of 1 mM auxin and *daf-2* RNAi treatment for 48 hours on the second day of adulthood. (D) Quantification of c) from n = 3 independent experiments. Error bars represent s.d. Two-sided *t*-test was used for statistical analysis. *: p < 0.05, **: p < 0.01, ***: p < 0.001. (E) Immunoblot analysis of DAF-2::degron animals showed no decrease of DAF-2 levels at high temperatures. Animals were raised at 15°C and put as L4 for 24 hours at the corresponding temperature. (F) A representative immunoblot analysis of “DAF-2::degron*”* animals after 1% glucose and 36 hours to 48 hours starvation on the second day of adulthood. (G) Quantification of f) from n = 3 independent experiments. Error bars represent s.d. Two-sided T-test was used for statistical analysis. *: p < 0.05, **: p < 0.01, ***: p < 0.001 (H) Immunoblot analysis of starved DAF-2::degron animals after eating HT115 (L4440) bacteria. Animals were raised on OP50 at 20°C and shifted as L4 to L4440 containing FUDR. After two days there were washed off and either frozen as control, or put on empty plates and harvested after 24 or 48 hours, respectively. (I) Immunoblot analysis of one-day-old adult DAF-2::degron animals treated with 1 mM auxin for the indicated time periods. (J) Quantification of e) from n = 3 independent experiments. Error bars represent s.d. One-sided t-test was used for statistical analysis. *: p < 0.05, **: p < 0.01, ***: p < 0.001. For (B-J): See Data Source File 1 and 2 for raw data, full blots, and statistics.

### Dietary changes modulate endogenous DAF-2/Insulin/IGF-1 receptor abundance

Next, we monitored endogenous DAF-2 protein levels under different environmental conditions, such as temperature and diet. Previously, Kimura and colleagues used DAF-2 antibody immunostaining of whole animals and reported that mutant DAF-2(*e1370*) protein is present at 15°C but barely detected at 25°C, whereas mutant DAF-2(*e1370*) protein in a *daf-16* null background or wild-type DAF-2 protein persists at both 15°C and 25°C (Kimura and Riddle, 2011). By contrast, upon 24 hours of starvation, the DAF-2 receptor is no longer detectable by using immunofluorescence in fixed *C. elegans* (Kimura and Riddle, 2011). Since the FOXO transcription factor *daf-16* is the transcriptional output of *daf-2* signaling (Ewald et al., 2018; Gems et al., 1998), these results suggest that DAF-2 protein levels may be autoregulated by insulin/IGF-1 signaling and might be influenced by temperature and food availability. We first asked whether our DAF-2::degron::3xFLAG tag allows quantification of endogenous DAF-2 levels. We observed comparable wild-type DAF-2::degron::3xFLAG levels across temperatures (15-28°C; Figure 1E), indicating that temperature does not influence DAF-2 levels in wild type. Intriguingly, however, we found that using FLAG-HRP antibodies to monitor protein levels, levels of DAF-2 protein almost completely disappeared after 36 to 48 hours of starvation (Figures 1F, 1G). In keeping with this result, well-fed animals, for which we added 1% glucose into the bacterial diet (OP50), increased the DAF-2 protein levels (Figures 1F, 1G). These effects were also influenced by the specific strain of *E. coli* used in each experiment: when *C. elegans* were fed HT1115 (L4440), DAF-2 levels did not decrease after 24 or 48 hours of starvation, suggesting that the nutritional composition of the animal’s diet prior to starvation influences DAF-2 stability (Figure 1H). Therefore, we conclude that different food sources and dietary cues control not only the secretion of insulin-like peptides to regulate DAF-2 activity (Pierce et al., 2001), but also DAF-2 receptor abundance directly.

### Auxin-induced degradation of degron-tagged Insulin/IGF-1 receptor

In *C. elegans*, cytosolic degron-tagged proteins are almost completely degraded after 30 minutes of auxin treatment (Zhang et al., 2015). However, the degradation of transmembrane proteins using AID *in vivo* has not been previously reported. We hypothesized that *C. elegans* might exhibit similar kinetics of degradation of a transmembrane protein following auxin treatment. In keeping with that hypothesis, after 30 minutes of 1 mM auxin treatment, we observed a dramatic decrease in transmembrane DAF-2 protein abundance (Figures 1I, 1J). Levels of DAF-2 were only slightly further reduced by continued auxin treatment, as indicated at 4 h and 24 h time points (Figures 1I, 1J). After 24 hours of 1 mM auxin treatment, we observed only a 40% total decrease in DAF-2 protein abundance, rather than a complete loss (Figures 1C, 1D). Similar kinetics in the degradation of DAF-2::degron::3xFLAG levels were confirmed using additional FLAG and degron antibodies (DataSource File 2). Taken together, these results suggest that our AID system allows for the partial, rapid degradation of the transmembrane DAF-2 receptor.

### Inactivation of DAF-2::degron by the AID inhibits downstream insulin/IGF-1 signaling

We wondered whether the reduction of DAF-2 levels seen with AID would result in reduced insulin/IGF-1 signaling that would have any physiological and functional consequences. Activation of DAF-2/insulin/IGF-1 receptor induces a downstream kinase cascade to phosphorylate the transcription factors DAF-16/FOXO and SKN-1/NRF, causing their retention in the cytoplasm (Figure 2A) (Ewald et al., 2015; Henderson and Johnson, 2001; Lin et al., 2001; Murphy, 2013; Ogg et al., 1997; Tullet et al., 2008). Genetic inhibition of *daf-2* results in less DAF-16 and SKN-1 phosphorylation and promotes nuclear translocation to induce the expression of target genes, such as *sod-3* (superoxide dismutase) and *gst-4* (glutathione S-transferase), respectively (Ewald et al., 2015; Henderson and Johnson, 2001; Lin et al., 2001; Murphy, 2013; Tullet et al., 2008). We found within 1 hour of 1 mM auxin treatment, most DAF-16::GFP translocated into the nuclei in DAF-2::degron animals (Figure 2B), with observable translocation already after 30 minutes (Supplementary Figure 2A). This DAF-16::GFP nuclear localization in DAF-2::degron animals was time-and auxin-concentration dependent and did not occur in DAF-16::GFP animals with wild-type DAF-2 (Figure 2B, Supplementary Figure 2B). Similarly, SKN-1- or DAF-16-target gene expression of *gst-4* or *sod-3* was only induced upon auxin treatment in DAF-2::degron animals (Figures 2C, 2D). Thus, insulin/IGF-1 signaling is reduced upon AID DAF-2 degradation.

**Figure 2:**
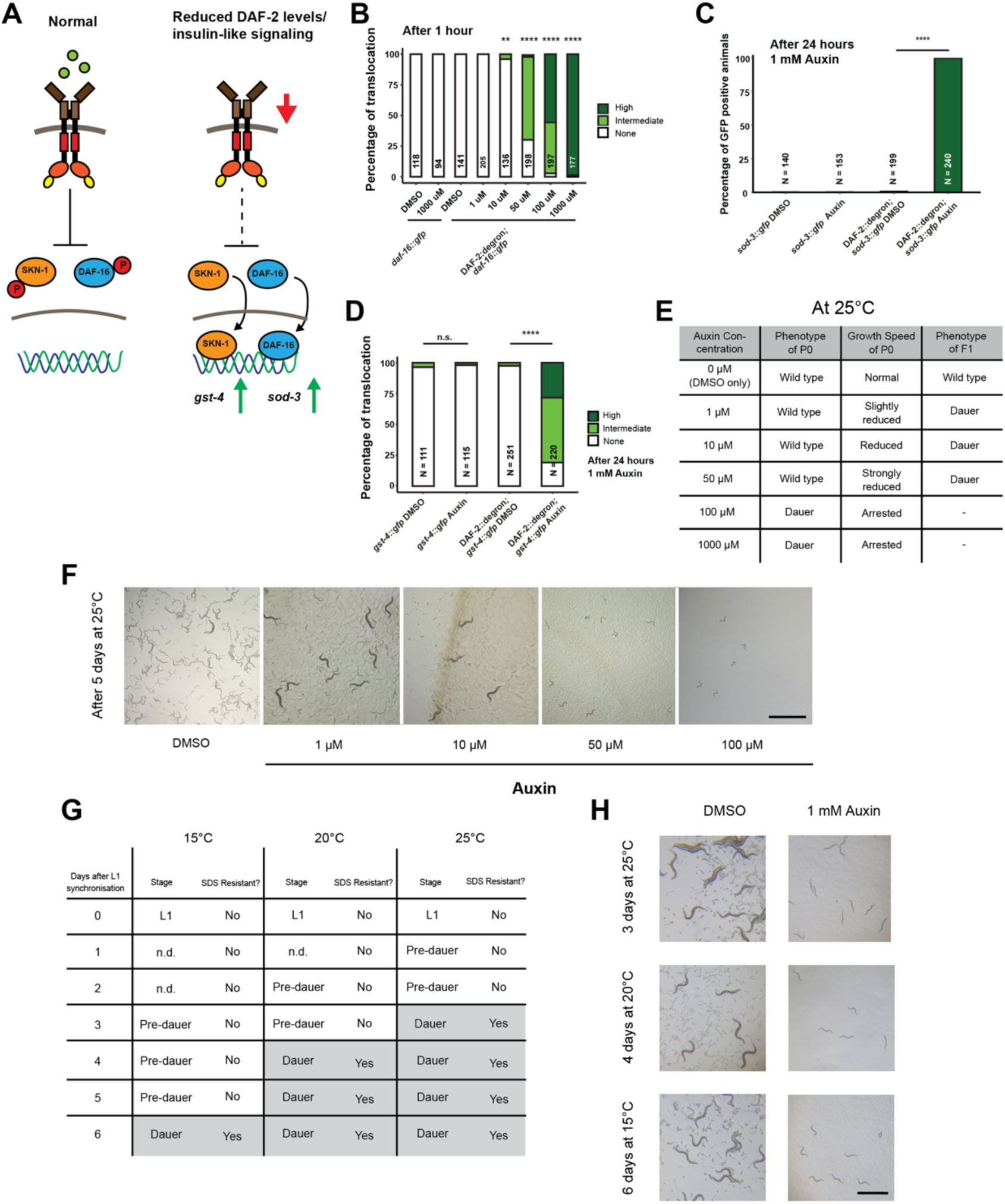
Inactivation of DAF-2::degron by the AID upregulated downstream reporters and caused dauer entry at any temperature. (A) A schematic illustration of the DAF-2 signaling pathway. DAF-2 phosphorylates the transcription factors DAF-16 and SKN-1 through a cascade of kinases and sequesters them to the cytosol. Lower DAF-2 levels or activity leads to de-phosphorylation and translocation of DAF-16 and SKN-1 to the nucleus and expression of targets genes like *sod-3* and *gst-4*, respectively. (B) One hour exposure to auxin led to nuclear translocation of DAF-16::GFP in a concentration-dependent manner in DAF-2::degron; *daf-16::gfp* animals, but not in *daf-16::gfp*. n > 94 from two (for *daf-16::gfp*) or three (for DAF-2::degron; *daf-16::gfp*) independent experiments. **: p < 0.01, ****: p < 0.0001. (C) Exposure to 1 mM auxin activated the reporter *sod-3::GFP* in DAF-2::degron*;sod-3::gfp* but not in animals that carry *sod-3::gfp* alone. Animals of the L4 stage were quantified after 24 hours. n > 140 from two (for *sod-3::gfp*) or four (for DAF-2::degron; *sod-3::gfp*) independent experiments. ****: p < 0.0001. (D) Exposure to 1 mM auxin activated the reporter *gst-4::GFP* in DAF-2::degron*;gst-4::gfp* but not in animals that carry *gst-4::gfp* alone. Animals of the L4 stage were quantified after 24 hours. n > 111 from two (for *gst-4::gfp*) or four (for DAF-2::degron; *gst-4::gfp*) independent experiments. ****: p < 0.0001. (E) Auxin treatment of DAF-2::degron affected development. Synchronized L1 put at low concentrations (1 to 50 μM auxin) show reduced growth speed, and their offspring enters dauer. At high concentrations (100 and 1000 μM), the L1 animals arrest and enter the dauer stage after a few days. (F) Representative pictures of growth impairment caused by auxin-mediated degradation of DAF-2 in DAF-2::degron animals at different concentrations. Bar = 1 mm. (G) Dauer entry of DAF-2::degron animals at 1 mM Auxin was temperature-independent, but the time needed for dauer entry is temporally scaled. To distinguish dauer animals from pre-dauer animals, they were treated for 15 minutes with 1% SDS. Only dauer animals survived SDS treatment. (H) Microscope pictures showed DAF-2::degron animals after dauer entry (right column) and control animals (left column) kept for the same amount of time at 15°C, 20°C, and 25°C, respectively. Control animals were on their second day of adulthood when the auxin-treated counterparts enter the dauer stage. Bar = 1 mm. For (B-D), see Data Source File 1 for raw data and statistics.

### AID of DAF-2::degron promotes dauer entry at any temperature

Reduced insulin/IGF-1 signaling during development promotes dauer entry. Dauer formation at 15°C has been observed for a variety of strong loss-of-function alleles, such as the class I alleles *e1369* and *m212*, the class II allele *e979*, the null alleles *m65, m646, m633,* and a variety of unclassified alleles discovered by Malone and Thomas (Gems et al., 1998; Kimura and Riddle, 2011; Malone and Thomas, 1994; Patel et al., 2008). For the commonly used reference alleles *e1368* and *e1370*, penetrant dauer formation only occurs at 25°C (Gems et al., 1998). By contrast, knockdown of *daf-2* by RNAi does not cause dauer formation at any temperature ^15,24,25^. We hypothesized that dauer formation would not happen because the decrease of DAF-2::degron levels after auxin treatment is only around 40%. To our surprise, synchronized L1 treated with 0.1 mM or 1 mM leads to dauer formation of all larvae at 25°C (Figures 2E, 2F). We observed a dose-dependent retardation of the developmental speed when using lower concentrations (1, 10, and 50 μM), but their offspring on plates containing auxin entered dauer (Figures 2E, 2F). The dependence on auxin concentration on dauer formation in L1 animals suggests an all-or-nothing threshold of DAF-2 receptor levels for the decision or commitment to dauer diapause. Even more surprising was that dauer formation was also observed at 15°C and 20°C with complete penetrance (Figures 2G, 2H). We found that dauer formation was temporally related to developmental speed at a given temperature: At 15°C, it took six days; at 20°C, it took four days; and at 25°C, it took three days to form dauers (Figure 2H). We verified that all auxin-induced DAF-2::degron dauers showed dauer-specific characteristics, such as SDS resistance, cessation of feeding, constricted pharynxes, and dauer-specific alae (Figure 2H, Supplementary Figures 2C-E), suggesting a complete dauer remodeling and transformation. Thus, AID of DAF-2 promotes complete dauer formation in a mechanism that is independent of temperature, but is dependent on DAF-2 protein abundance.

### Dauer commitment at mid-larval stage one upon DAF-2 degradation

Wild-type animals enter the pre-dauer L2d stage, where they keep monitoring their environment before completely committing to dauer formation (Hu, 2007; Karp, 2018). However, previous temperature-shifting experiments (from 15°C to 25°C) with *daf-2* mutants suggested a dauer decision time-window from L1 to L2 stage before the L2d stage (Swanson and Riddle, 1981). The AID allows for precise temporal degradation of DAF-2. We pinpointed the dauer entry decision to mid-L1 by shifting synchronized L1 at different time points to plates containing 1 mM auxin and by counting the number of germ cell precursors to determine the exact age of the population at the time point when roughly 50% of the population committed to dauer stage (Supplementary Figures 3A-C). We found that when DAF-2 levels are below a given threshold at mid-L1 stage, the animals commit to dauer formation irrespective of the environment during L2d or later stages.

### Auxin-induced degradation of DAF-2 resembles a non-conditional and severe loss-of-function allele

The FOXO transcription factor DAF-16 is required for dauer formation in *daf-2* mutants. We crossed DAF-2::degron with DAF-16::degron (Hobert, 2019) and found that *daf-16* was required for dauer formation and developmental speed alterations after DAF-2::degron depletion (Supplementary Figure 3D). Previously reports suggest that many *daf-2* alleles show low to severe penetrance of embryonic lethality and L1 arrest at higher temperatures (Collins et al., 2008; Ewald et al., 2016; Gems et al., 1998; Patel et al., 2008). Although the constitutive dauer formation of the proposed null allele *daf-2(m65)* is suppressed by *daf-16* null mutations, the embryonic lethality and L1 arrest are *not daf-16* dependent (Patel et al., 2008). Surprisingly, we observed no embryonic lethality or L1 arrest in the progeny of animals placed animals on 1 mM auxin as L4s. Similar results were seen using either the DAF-2::degron or DAF-2::degron; germline TIR1 strains. Taken together, these results suggest that, although there is still DAF-2::degron::3xFLAG detected with the FLAG-HRP antibody after 24 hours of auxin treatment, the persistent pool of DAF-2 receptor is insufficient to provide the wild-type functions. Higher concentrations of auxin treatment lead to toxicity in both wild type and DAF-2::degron animals (Supplementary Figure 3E). Inactivation of DAF-2 by the AID is 100% penetrant for dauer formation at any temperature. Still, the absence of embryonic lethality or L1 arrest at 1 mM auxin suggests that DAF-2::degron functionally is more similar to a non-conditional and severe loss-of-function mutation than a null allele.

### Enhanced lifespan extension by AID of DAF-2 in adult animals

Given the strong phenotypic effects of DAF-2 AID on animal development, we next explored whether DAF-2 degradation by AID could affect the function of adult animals. Previous studies indicate that reducing IIS either by *daf-2* RNAi knockdown or in genetic mutants increases lifespan at any temperature (15-25°C) (Ewald et al., 2018; 2015; Gems et al., 1998). We hypothesized that AID-dependent degradation of DAF-2 would have similar effects on the lifespan of animals. We found that auxin supplementation of DAF-2::degron animals, starting from L4, resulted in a 70-135% lifespan extension (Figure 3A; Supplementary Table 1). DAF-2 degradation using 1 mM auxin surpassed the longevity of commonly used *daf-2(e1368)* and *daf-2(e1370)* mutants (Figure 3A, Supplementary Table 1). By contrast, auxin-treatment at 0.1 mM or 1 mM concentration had little or no effect on wild-type lifespan (Figure 3A; Supplementary Table 1). These results suggest that auxin-induced degradation of *daf-2* is a powerful tool to promote longevity.

**Figure 3.**
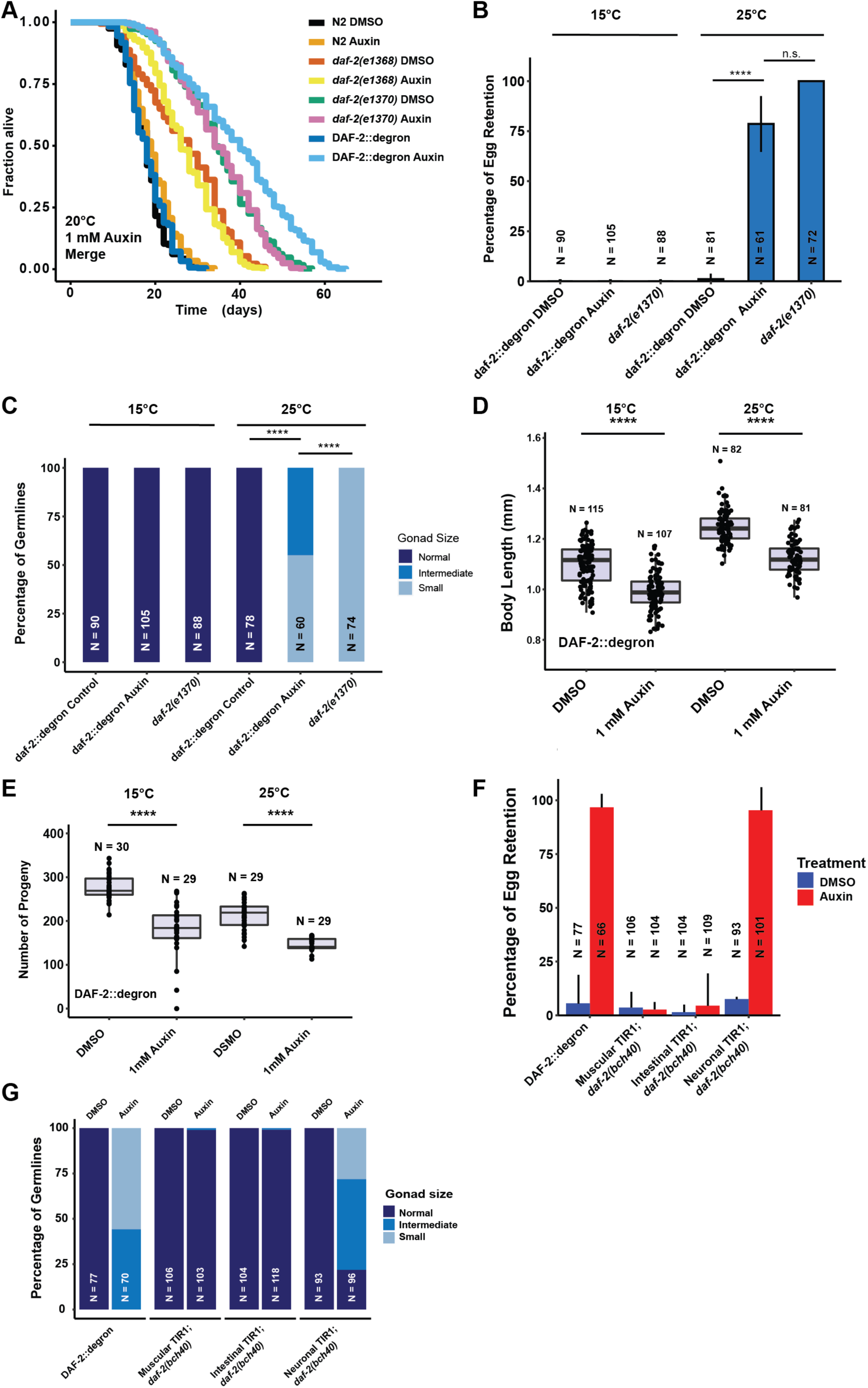
Depletion of DAF-2::degron causes dauer-associated phenotypes at 15°C. (A) Auxin treatment extended the lifespan of DAF-2::degron animals. Animals were shifted as L4 to plates containing DMSO or 1 mM auxin at 20°C. (B) 1 mM auxin treatment leads to the “dauer-associated” phenotype egg retention in DAF-2::degron. Animals were raised at 15°C and shifted as L4 to 25°C or kept at 15°C. Two days later, at 25°C and three days later at 15°C, the animals were checked for egg retention. The experiment was performed 3 independent times. Error bar represents s.d. ****: p < 0.0001 (C) Gonads were shrunk after 1 mM auxin treatment in DAF-2::degron animals. Animals were raised at 15°C and shifted as L4 to 25°C or kept at 15°C. Two days later at 25°C and three days later at 15°C, the animals were checked for gonad size. The experiment was performed 3 independent times. ****: p < 0.0001 (D) 1 mM auxin treatment decreased the body size of DAF-2::degron animals at 15°C and 25°C. Animals were raised at 15°C and shifted to 1 mM auxin or DMSO plates at the L4 stage. The experiment was performed 3 independent times. ****: p < 0.0001 (E) 1 mM auxin treatment of DAF-2::degron resulted in a smaller brood size at 15°C and 25°C. Animals were shifted to 1 mM auxin or DMSO plates at the L4 stage and either kept at 15°C or moved to 25°C. The experiment was performed 3 independent times. ****: p < 0.0001 (F) Tissue-specific depletion of DAF-2 in neurons caused egg retention phenotype. Animals were raised at 15°C and shifted from L4 to 25°C. Two days later, the animals were checked for egg retention. The experiment was performed 3 independent times. Error bar represents s.d. ****: p < 0.0001 (G) Gonads were shrunk after neuronal depletion of DAF-2. Animals were raised at 15°C and shifted from L4 to 25°C. Two days later, the animals were checked for gonad size. The experiment was performed 3 independent times. ****: p < 0.0001 For (A-G), see Data Source File 1 for raw data and Supplementary Table 1 for statistics and additional trials.

### Manifestation of some dauer-associated phenotypes during adulthood at 15°C without passing through L2D

Although reducing *daf-2* function causes beneficial increases in longevity and stress resistance, it causes residual detrimental phenotypes in adult animals that resemble the behavioral and morphologic changes expected in a dauer state (Ewald et al., 2018; Gems et al., 1998). In class II *daf-2* alleles, dauer-associated phenotypes, such as small gonads, reduced brood size, reduced motility, and reduced brood size, manifest only at 25°C during adulthood, but not at lower temperatures (Ewald et al., 2018; Gems et al., 1998). To determine whether DAF-2::degron AID animals display dauer-associated phenotypes, we quantified these dauer-like characteristics at 15°C and 25°C. In placing L4 DAF-2::degron animals on auxin and at 25°C, we observed similar levels of egg retention and effects on gonad size as was seen in the *daf-2(e1370)* class II allele. Strikingly, these effects were temperature-dependent, as DAF-2::degron animals did not retain eggs or have reduced gonad sizes at 15°C (Figure 3B, Supplementary Figure 4A). Similarly, DAF-2::degron animals on auxin exhibited germline shrinkage at 25°C, albeit to a lesser degree than the *daf-2(e1370)* class II allele (Figure 3C, Supplementary Figure 4B). This phenotype was also absent at 15°C, suggesting that egg retention and germline-shrinkage reminiscent of dauer-associated remodeling phenotypes are temperature-sensitive traits.

Another known dauer-associated phenotype at 25°C is the quiescence or immobility of class II *daf-2(e1370)* mutants (Ewald et al., 2015; Gems et al., 1998). We did not observe any immobility of auxin-treated DAF-2::degron animals at 25°C or during lifespan assays at 20°C (Supplementary Video 1, Supplementary Table 2). Although the effects on body size of *daf-2(e1370)* class II allele are temperature dependent, presenting at 25°C but not at 15°C (Ewald et al., 2015; Gems et al., 1998; McCulloch and Gems, 2003) (Supplementary Figure 4C), auxin-induced degradation of DAF-2 starting from L4 shortened body size of 2-days-old adults at both temperatures (Figure 3D, Supplementary Figure 4D). Similarly, while *daf-2(e1370)* mutants only exhibit reduced brood sizes at higher temperatures (Ewald et al., 2015; Gems et al., 1998); AID of DAF-2::degron starting from L4 reduced brood size at both 15°C and 25°C (Figure 3E). This suggests that lower body and brood size manifest as non-conditional traits, in keeping with insulin/IGF-1’s role as an essential gene for these functions. In summary, these results suggest that some dauer-associated phenotypes or pathologies can be induced during adulthood independent of temperature and that passing through L2d is not required for dauer-associated phenotypes in adult animals.

### Tissue-specific DAF-2 degradation reveals neuronal regulation of egg retention and germline remodeling

The pleiotropic effects of DAF-2 have been ascribed to tissue-specific effects of DAF-2 function. DAF-2 protein levels are predominantly found in the nervous system and intestine, and to a lesser extent in the hypodermis (Kimura and Riddle, 2011), while *daf-2* mRNA expression has also been detected in the germline (Han et al., 2017; Lopez et al., 2013). Importantly, dauer formation in *daf-2(e1370)* can be restored by expressing wild-type DAF-2 only in neurons (Wolkow et al., 2000). We hypothesized that select tissues might drive these dauer-associated phenotypes. To test this, we expressed TIR1 specifically in muscles, neurons, and intestine driven by the *myo-3, rab-3,* and *vha-6* promoters, respectively (Materials and Methods, Supplementary Table 2). TIR1 expressed from any of these three tissue-specific promoters did not result in reduced body size (Supplementary Figure 4E). To validate that the neuronal TIR1 was functional, we crossed neuronal TIR1 into *daf-16(ot853* [*daf-16*::linker::mNG::3xFLAG::AID]) (Hobert, 2019) and observed that DAF-16::mNG was selectively degraded in neurons upon auxin treatment (Supplementary Figure 5A). We found that depletion of DAF-2 in neurons caused egg retention and germline shrinkage (Figures 3F, 3E). Interestingly, the germline shrinkage and egg retention phenotypes were temperature-dependent, suggesting some interaction of temperature and neuronal DAF-2 abundance. Thus, some *daf-2*-phenotypes are initiated from a single tissue, whereas others might be due to a multiple tissue interplay.

### Tissue-specific AID reveals different requirements for DAF-2 in neurons and intestine for longevity and oxidative stress resistance

Previously, transgenic expression of wild-type copies of DAF-2 in neurons or intestine was shown to partially suppress the longevity of *daf-2(e1370)* mutants at 25°C (Wolkow et al., 2000). Taking advantage of our unique AID system, we wanted to ask whether the degradation of DAF-2 in a single tissue would be sufficient to induce longevity. We found that either neuronal or intestinal depletion of DAF-2 alone was sufficient to extend lifespan, although not to the extent as when DAF-2 is degraded in all tissues (Figure 4A; Supplementary Table 1). Therefore, we asked whether tissue-specific DAF-2 degradation was also sufficient for stress resistance seen in *daf-2* mutant animals (Ewald et al., 2018; Gems et al., 1998). Auxin-induced degradation of DAF-2 in all tissues resulted in increased oxidative stress resistance, comparable to previous for *daf-2(e1370)* mutants (Figure 4B). We observed improved oxidative stress resistance when we depleted DAF-2 in the intestine, but not in neurons (Figures 4C, 4D, Supplementary Figures 5b, 5C). This implies that DAF-2 in different tissues orchestrates systemic signaling for organismal lifespan and stress resistance. Furthermore, reducing DAF-2 levels, specifically in the intestine, promotes longevity and stress resistance without causing dauer-associated phenotypes.

**Figure 4.**
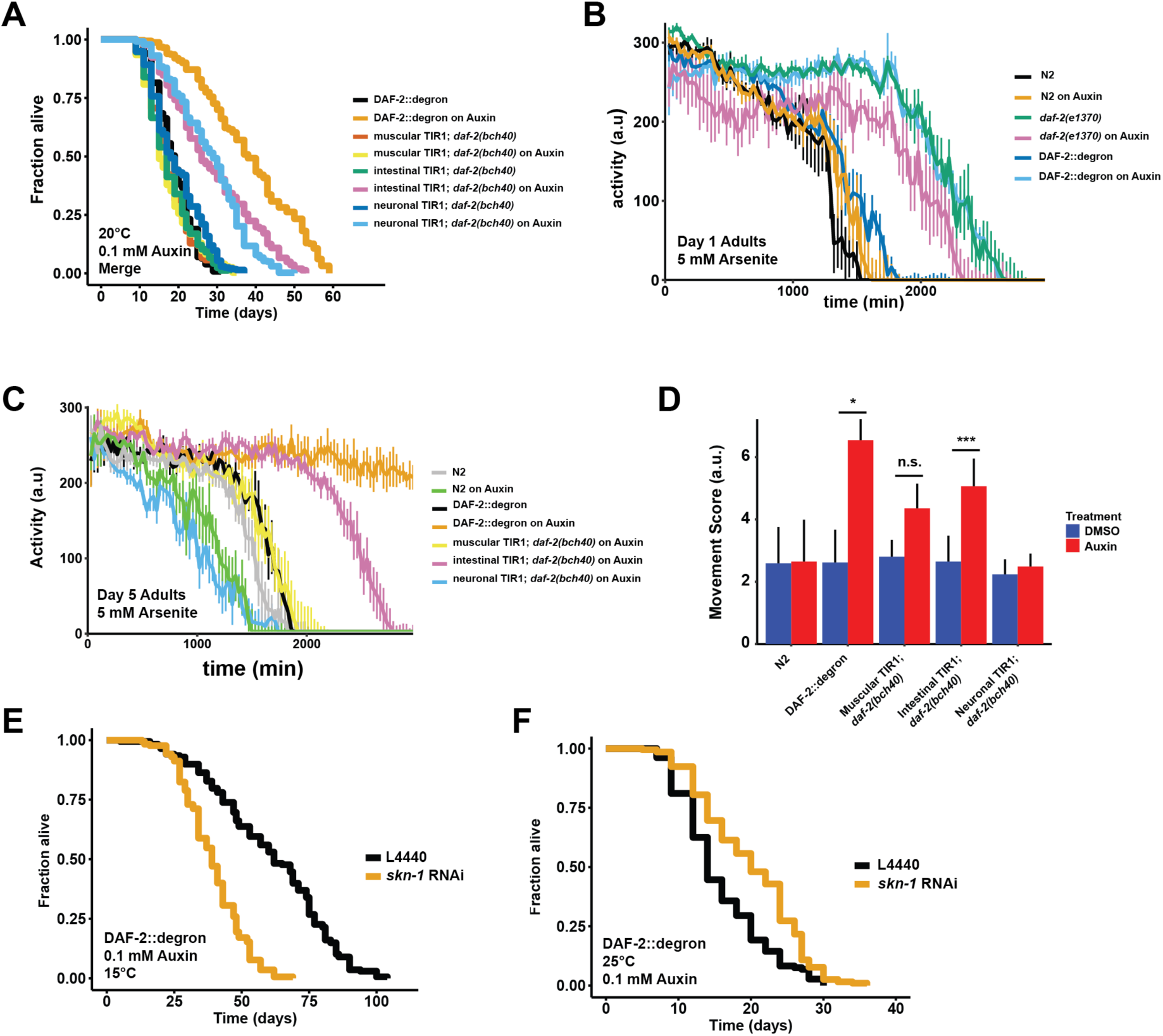
Tissue-specific effects after DAF-2 depletion on lifespan extension and reactive oxygen species resistance. (A) Auxin treatment extended lifespan in aged DAF-2::degron animals. Animals were shifted from DMSO plates to plates containing 1 mM Auxin on the indicated day. (B) Auxin-mediated depletion of DAF-2 enhanced oxidative stress resistance to a similar extent of *daf-2(e1370)* animals. Animals were shifted as L4 for 24 hours on plates containing DMSO or 1 mM auxin at 20°C. (C) Intestinal knockdown of DAF-2 enhanced oxidative stress resistance. Animals were shifted to DMSO or 1 mM auxin plates at L4 stage and kept until day 5 of adulthood at 20°C. (D) Quantification of g) from 3 independently performed experiments. Error bar represents s.d. *: p < 0.05, ***: p < 0.001. (E) *skn-1* was necessary for full lifespan extension in DAF-2 depleted animals at 15°C. Animals were kept on 1 mM auxin. p < 0.0001 (F) Knockdown of *skn-1* extended the mean lifespan of DAF-2 depleted animals at 25°C. Animals were raised at 15°C and shifted to 1 mM auxin at 25°C. p < 0.0001 For (A-F), see Data Source File 1 and 3 for raw data and Supplementary Table 1 for statistics and additional trials.

### *skn-1* works in a temperature-sensitive manner but independently of dauer-associated reprogramming

We have previously shown that the transcription factor SKN-1/NRF1,2,3 is localized in the nucleus at 15°C or 25°C in *daf-2(e1370)* mutants (Ewald et al., 2015), suggesting SKN-1 activation occurs under reduced insulin/IGF-1 receptor signaling conditions (Tullet et al., 2008). Intriguingly, *skn-1* activity is necessary for full lifespan extension of *daf-2(e1370)* mutants only at 15°C but not at 25°C (Ewald et al., 2015). We hypothesized that *skn-1* requirements for lifespan extension become masked when dauer-associated reprogramming conditions are triggered at the higher temperatures (Ewald et al., 2018). Because auxin-treated DAF-2::degron AID animals exhibit dauer-associated traits during adulthood at 15°C, we asked whether the lifespan extension caused by DAF-2::degron AID upon auxin treatment at 15°C is *skn-1*-independent. We found that the lifespan extension in DAF-2::degron animals fully required *skn-1* at 15°C. Surprisingly, however, a loss of *skn-1* extended the median lifespan of DAF-2::degron animals at 25°C (Figures 4E, 4F; Supplementary Table 1). This suggests that *skn-1* may function independently from dauer-associated reprogramming pathways at 15°C. Furthermore, the increased longevity seen in DAF-2::degron animals may result from a differential transcriptional program at higher temperatures compared to lower temperatures.

### Late-life application of AID of DAF-2 increases lifespan

Finally, we asked whether it would be possible to promote longevity in extreme geriatric animals by depleting DAF-2 by AID. Previous studies using RNAi indicated that reduced *daf-2* expression extended lifespan when started at day 6 of adulthood but not later (Dillin et al., 2002), raising the question of whether *daf-2*-longevity induction is possible beyond the reproductive period (day 1-8 of adulthood). To address this, we maintained DAF-2::degron animals on control plates and shifted them to 1 mM auxin containing plates at day 0 (L4) up to day-20 of adulthood (Figure 5A). We found that shifting the animals past the reproductive period at day 10 and day 12 still led to an increase in lifespan by 48-72% and 49-57%, respectively (Figure 5A, Supplementary Table 1). Since transferring old *C. elegans* to culturing plates without bacterial food can also increase lifespan past reproduction (Smith et al., 2008), we decided to top-coat lifespan plates with auxin late in life. We observed lifespan extension of animals by supplementing auxin very late during lifespan at day 25 of adulthood, at a time at which already approximately 50% of the population had died (Figures 5B-D, Supplementary Table 1). This demonstrates that reducing insulin/IGF-1 receptor signaling is feasible in geriatric *C. elegans.* Thus, our use of AID to selectively degrade the DAF-2 protein suggests that targeting the *daf-2* signaling pathway late in life is sufficient to extend lifespan.

**Figure 5.**
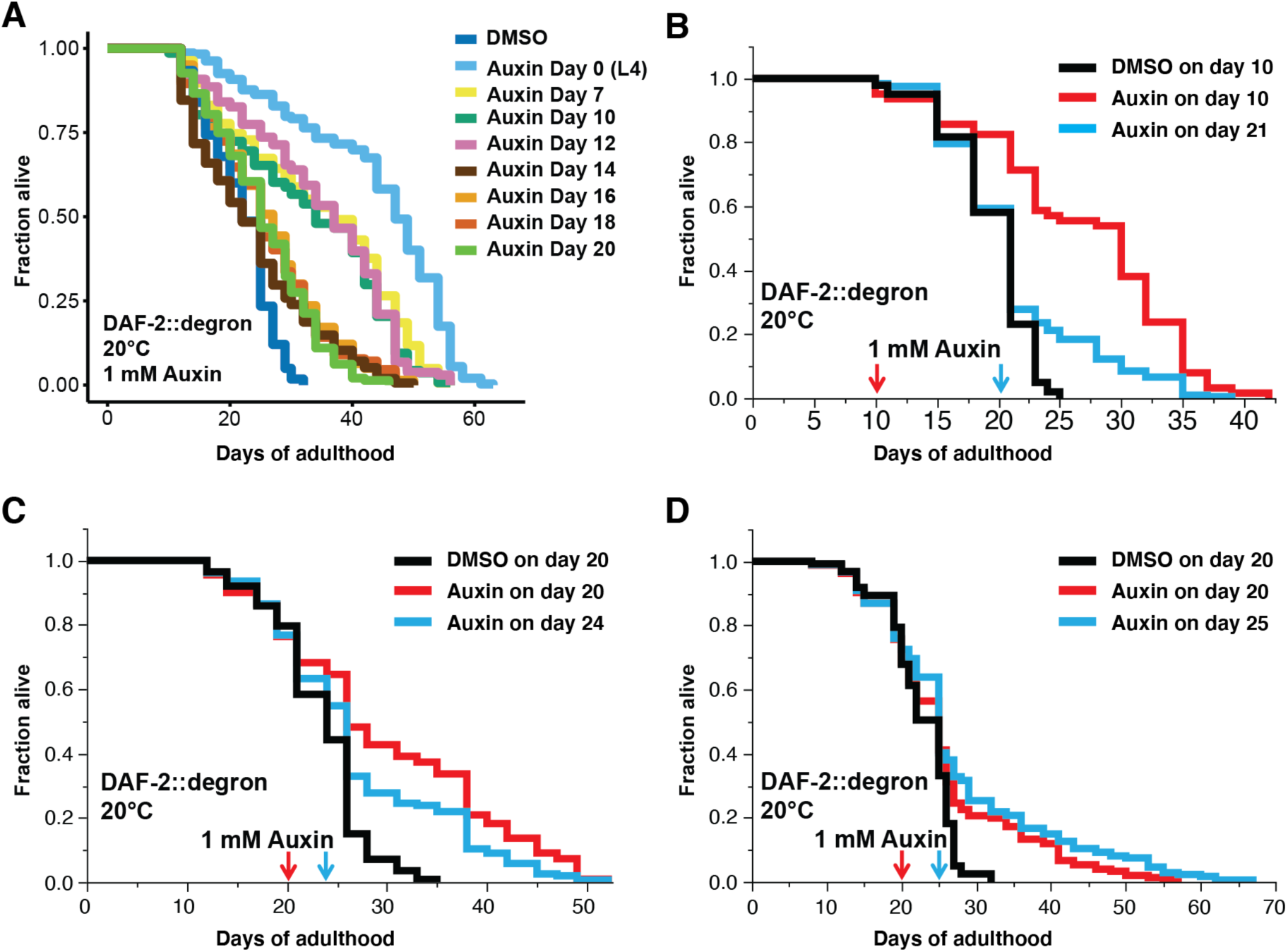
Late-life DAF-2 depletion extended lifespan. (A) Auxin treatment extended lifespan in aged DAF-2::degron animals. Animals were shifted from DMSO plates to plates containing 1 mM Auxin at the indicated day. (B-C) Top-coating plates with auxin to a final concentration of 1 mM auxin at the indicated days extended the lifespan of DAF-2::degron animals during aging. For (A-D), see Supplementary Table 1 and Data Source File 1 for raw data, statistics, and additional independent trials.

## Discussion

For longevity interventions to be efficient without causing undesired side effects, the time point of treatment must be chosen carefully. This is especially important for pathways such as the insulin/IGF-1 pathways that are essential for growth and development (Kenyon, 2010). Although the importance of DAF-2 in regulating lifespan is well-established, the consequences of late-in-life inhibition remained unknown.

Here we demonstrate that late-life degradation of *daf-2* extends lifespan. In this work, we have effectively engineered a degron tag into the endogenous *daf-2* locus using CRISPR, representing the first report of auxin-induced degradation (AID) of a transmembrane receptor *in vivo*. DAF-2 receptor levels are efficiently degraded via AID. AID of DAF-2 is functional at three distinct life stages, affecting dauer formation specifically during development, longevity, stress resistance, growth-related phenotypes during early adulthood, and finally longevity during post-reproductive geriatric stages.

It is well-established that DAF-2/insulin/IGF-1 receptor signaling connects nutrient levels to growth and development (Murphy, 2013). This is attributed to insulin-like peptides binding DAF-2 and activating a downstream phosphorylation kinase cascade that adapts metabolism (Murphy, 2013). Surprisingly, we find that DAF-2 receptor abundance is linked to nutrient availability and dietary content. Starving *C. elegans* decreases DAF-2 receptor abundance consistent with previous *in-situ* antibody staining (Kimura and Riddle, 2011), whereas mimicking a high-energy diet by adding glucose to the bacterial food source increases DAF-2 receptor abundance. This suggests an additional layer of regulating insulin/IGF-1 signaling by connecting food cues to adapt metabolism via DAF-2 receptor levels, potentially as an internal representation of the environment.

Environmental conditions are carefully monitored by developing *C. elegans*. First, larval stage (L1) wild-type *C. elegans* that are in food-deprived environments, that are surrounded by elevated temperatures (>27°C), and/or that are in high population densities into a pre-dauer L2d stage, where they continue monitoring their environment prior to completely committing and molting into dauers (Hu, 2007; Karp, 2018). However, previous temperature-shift experiments (from 15°C to 25°C) with *daf-2* mutants have indicated the existence of a “dauer decision time-window” between the L1 and L2 stage (Swanson and Riddle, 1981). By manipulating DAF-2 levels directly using AID, we found that AID-degradation of DAF-2 during a narrow time period in the mid-L1 stage is sufficient to induce dauer formation, suggesting that the decision to enter dauer relies upon an all-or-nothing threshold of DAF-2 protein levels. This decision is uncoupled from temperature or food abundance and occurs irrespective of the conditions present later during L2 or L2d-predauer. Thus, absolute DAF-2 protein abundance appears to be the key (if not sole) factor in the animal’s decision to enter into the dauer state during early development.

Although IIS is reduced in *daf-2(e1370* or *e1368)* mutants at lower temperatures (Ewald et al., 2018; 2015; Gems et al., 1998), dauer larvae are formed only when these *daf-2* mutants are exposed to higher temperatures. The observation that mutant DAF-2 protein (Kimura and Riddle, 2011) but not wild-type DAF-2 protein (Figure 5) is lost at more elevated temperatures suggests a model in which the DAF-2 mutant protein becomes unstable with increasing temperatures and might be subsequently targeted for degradation. This hypothesis helps to explain why strong class II *daf-2* mutants, such as *daf-2(e979)* and *daf-2(e1391)*, which have much lower DAF-2 protein levels at 15°C and 25°C compared to other *daf-2* mutants (Tawo et al., 2017), exhibit higher propensities toward dauer formation at any temperature (Gems et al., 1998). Thus, the severity of classical *daf-2* mutant alleles in regards to dauer formation and adulthood dauer traits might also be linked to DAF-2 receptor abundance.

At all temperatures, *daf-2* mutants go through an alternative pre-dauer stage (L2d) even if they later develop into gravid adults (Karp, 2018; Ruaud et al., 2011). This suggests that *daf-2* mutants may instead commit to an alternative developmental path that results in reprogrammed physiology carried over into adulthood. This is evident in adult *daf-2* class II mutants at higher temperatures, as they still exhibit significant levels of “dauer-associated phenotypes” (Ewald et al., 2018). In these animals, a clear remodeling of the body and internal organs, including constriction of the pharynx and shrinkage of the germline, remains present, along with an altered neuronal morphology and electrical synapse connectome that drives behavioral changes such as quiescence, diminished foraging behavior, and altered egg-laying programs ^12,15,19-23,36^. Since adult-specific treatment with *daf-2* RNAi increases lifespan without eliciting these dauer-associated traits (Ewald et al., 2015), this has suggested that these dauer traits of *daf-2* mutants are remnants of the animals’ time in the “alternative” L2d developmental stage. In this study, however, we show that L4-specific AID of the DAF-2::degron results in non-conditional reduction of body size and brood size, whereas egg retention and germline shrinkage only occurs at higher temperatures. This indicates that the non-conditional dauer traits are not a residual effect of the L2d developmental program but instead are side-effects caused by reducing the functions of DAF-2 related to the regulation of growth.

Additional temperature-sensitive dauer traits of egg-retention and gonad shrinkage are mediated by loss of *daf-2* in neurons only at higher temperatures. Why these traits only manifest at higher temperature remains unclear. One explanation may be that DAF-2 levels are reduced in temperature-sensing neurons, which then elicits a systemic effect that drives germline shrinkage and egg-retention. Neurons are refractory to RNAi, suggesting that *daf-2(RNAi)* effects work through other tissues than neurons to extend lifespan. Furthermore, treating Class I mutants *daf-2(e1368)* with *daf-2(RNAi)* doubles their longevity without causing adult dauer traits (Arantes-Oliveira et al., 2003), suggesting that *daf-2(RNAi)* would lower DAF-2 receptor levels in other tissues than neurons for this additive longevity effect. Consistent with neuronal regulation of these traits is that *daf-2* RNAi applied to wild type does not result in dauers but when applied to neuronal-hypersensitive RNAi *C. elegans* strains results in dauer formation (Dillin et al., 2002; Ewald et al., 2015; S. Kennedy et al., 2004). We find that DAF-2 degradation in neurons or intestine increases lifespan. Given that reducing DAF-2 in neurons results in dauer traits at higher temperatures, one might target intestinal DAF-2 for degradation to uncouple from any dauer traits. Yet, DAF-2 is essential for growth. We find the best time point for DAF-2 inhibition is rather late in life to by-pass these undesired side-effects to promote longevity.

We find that as late as day-25 of adulthood, when about 50% of the population has died, AID of DAF-2 is sufficient to increase lifespan. The only application that was able to also increase the lifespan that late was transferring old *C. elegans* to culturing plates without bacterial food (Smith et al., 2008). But *C. elegans* do not feed after reaching mid-life (Collins et al., 2008; Ewald et al., 2016), suggesting it is not the intake of calories that promotes longevity. Could it be that the old *C. elegans* smell or sense the absence of food and thereby reduce DAF-2 levels to promote longevity? It would be interesting to determine if late-life bacterial deprivation works synergistically to DAF-2 AID or not in future studies.

In mammals, mid-life administration of IGF-1 receptor monoclonal antibodies to 78-weeks old mice (a time point at which all mice are still alive and 6-weeks before first mice start to die) is sufficient to increase their lifespan and improves their healthspan (Mao et al., 2018). Other parallels between mammals and nematodes are the tissues from which the insulin/IGF-1 receptor induces longevity. Brain-specific heterozygous IGF-1 receptor knockout mice are long-lived (Kappeler et al., 2008), whereas adipose-specific insulin receptor knockout results in longevity (Blüher et al., 2003). Since *daf-2* is considered the common ancestor gene to both insulin receptor and IGF-1 receptor (Kimura et al., 1997), it is interesting that AID of DAF-2::degron in neurons or in the major fat-storage tissue intestine was sufficient to increase the lifespan of *C. elegans*.

Although it is known that neurons and intestine are important for food perception and regulation of food intake, the effects of food perception or intake on insulin receptor and IGF-1 receptor levels is less understood. Starving rats for three days increases the abundance of insulin-bound insulin receptors (KOOPMANS et al., 1995), maybe to allow more glucose uptake. It is unknown whether prolonged starvation would lead to lower basal insulin/IGF-1 receptor levels. However, genetic alterations of the IGF-1 receptor levels are associated with altered lifespan. For instance, heterozygous IGF-1 receptor knockout mice, which have lower IGF-1 receptor levels, have an increased lifespan (Holzenberger et al., 2003; Kappeler et al., 2008; Xu et al., 2014). Overexpression of the short isoform of p53 (p44) increases IGF-1 receptor levels and shortens the lifespan of mice (Maier et al., 2004). Furthermore, administration of recombinant human IGF-1 increases IGF-1 receptor abundance in murine embryonic cells (Maier et al., 2004). Food components themselves can affect insulin receptor levels. For instance, palmitate activates PPAR*α* to induce miR-15b that binds insulin receptor mRNA for degradation (Li et al., 2019). Whether food components can regulate insulin/IGF-1 receptor levels via E3 ligase-mediated degradation in mice would be an exciting aspect for future research.

There is some evidence that E3 ligases have important physiological functions in regulating insulin/IGF-1 receptor abundance. For instance, endogenous E3 ligase CHIP regulates insulin/IGF-1 receptor levels in *C. elegans*, *Drosophila*, and human cell cultures (Tawo et al., 2017). In mice, the muscle-specific mitsugumin 53 (MG53) E3 ligase targets the insulin receptor for degradation (Song et al., 2013). High-fat diet results in the reduction of insulin receptor levels (Li et al., 2019) via higher MG53-mediated degradation (Song et al., 2013). MG53 is upregulated under a high-fat diet in mice, and MG53-/- deficient mice are protected from high-fat diet-induced obesity, insulin resistance, and other metabolic syndrome associated phenotypes (Song et al., 2013). Furthermore, another E3 ligase MARCH1, is overexpressed in obese humans and targets the insulin receptor for ubiquitin-mediated degradation (Nagarajan et al., 2016). Taken together, this suggests that food abundance controls mammalian insulin receptor levels via E3 ligase targeted degradation. Although in *C. elegans,* we found the opposite changes in DAF-2 receptor levels, which were reduced upon starvation and increased upon high glucose feeding, it suggests that nutritional cues regulate insulin/IGF-1 receptor levels via a variety of mechanisms, including ubiquitination and proteasomal degradation, across species.

In summary, we have demonstrated that late-life interventions can increase lifespan. We have established that auxin-induced degradation is suitable for targeting transmembrane receptors for non-invasive manipulations during developmental and longevity *in-vivo* studies. Using the AID of DAF-2, we reconciled a longstanding question by providing evidence that dauer-traits are not a spill-over of reprogrammed physiology from developing L2d pre-dauers. Instead, the essential growth-related functions of DAF-2 are causing deficits when applied during development or growth phases. We have shown that tissue-specific or interventions beyond reproduction or growth extend lifespan without pathology or deficits. Degradation of DAF-2/insulin/IGF-1 receptor might not be an artificial intervention since DAF-2/insulin/IGF-1 receptor abundance is read-out to adapt metabolism to the environment and food status. E3 ligases might play an important role in regulating DAF-2/insulin/IGF-1 receptor levels. Dissecting intrinsic DAF-2/Insulin/IGF-1 receptor abundance in response to nutritional cues may impact our understanding of nutrient-sensing in promoting health during old age.

---

**Supplementary Figure 1.**
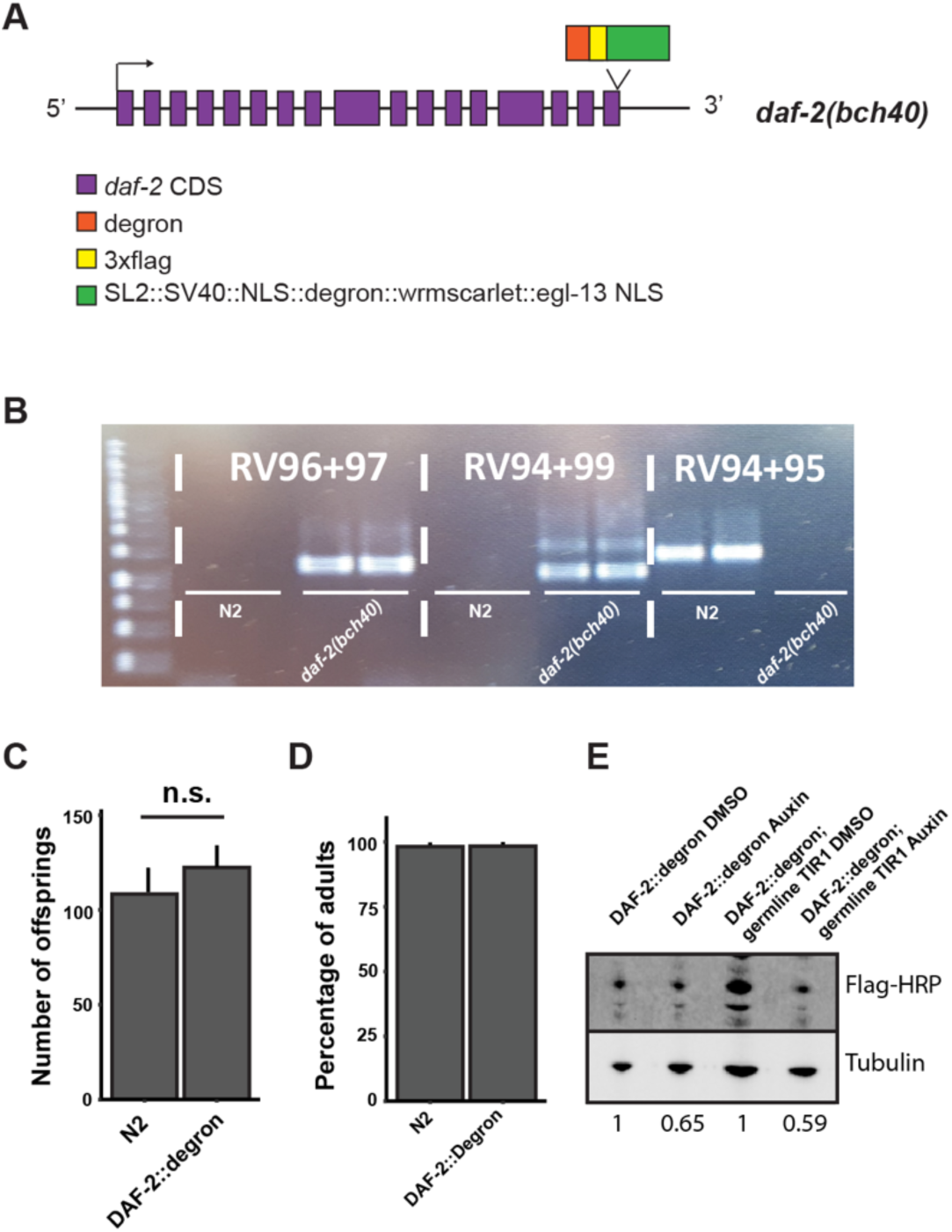
Degron-tagged DAF-2 is functional and susceptible to auxin-mediated degradation. (A) Schematic illustration of the *daf-2* locus of *daf-2(bch40* [*daf-2::degron::3xFLAG::STOP::SL2::SV40::degron::wrmScarlet::egl-13NLS*]) animals. Expression of the operon was not observed. Either it is not functional, or the levels are too low to be detected by confocal microscopy. (B) PCR of wild type (N2) and *daf-2(bch40)*. Primer pair RV96 and RV97 span the 3’ of the insert, RV94 and RV99 span the 5’ of the insert, and RV94 and RV95 span the end of the gene of the wild type. The PCR product of RV96 and RV97 and RV94 and RV99 have been sequenced to verify the construct’s correct insertion. (C) Comparison of progeny number of wild type (N2) and DAF-2::degron from n = 3 independent experiments. Error bar represents s.d. (D) Comparison of the developmental speed of wild type (N2) and DAF-2::degron from n = 3 independent experiments. Error bar represents s.d. (E) Immunoblot analysis of DAF-2::degron and DAF-2::degron; germline TIR1 (*eft-3p::TIR1::mRuby::unc-54* 3’UTR; *daf-2(bch40); sun-1p::TIR1::mRuby::sun-*1 3’UTR) on DMSO and 1 mM auxin. The numbers below indicate the normalized value to the respective control on DMSO. For (C-E), see Data Source File 1 and 2 for raw data and statistics.

**Supplementary Figure 2:**
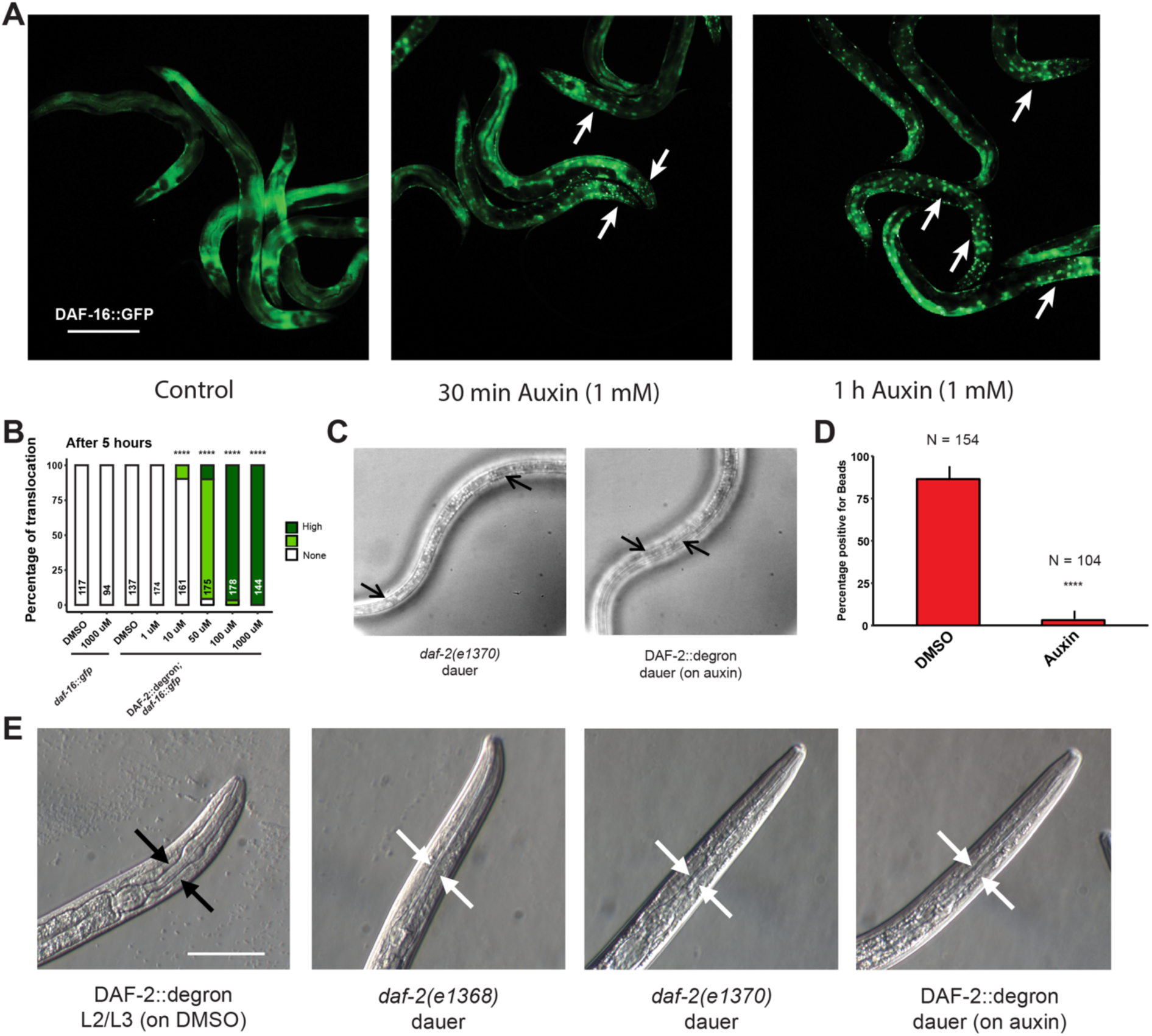
Inactivation of DAF-2::degron by the AID upregulated downstream reporters and caused dauer entry at any temperature. (A) Translocation of DAF-16::GFP in *“DAF-2::degron”; daf-16::gfp* animals was visible after 30 minutes and 1 hour of 1 mM auxin treatment. Bar = 100 uM. (B) Five hour exposure to auxin led to further nuclear translocation of DAF-16::GFP in a concentration-dependent manner in DAF-2::degron; *daf-16::gfp*, but not in *daf-16::gfp*. n > 94 from two (for *daf-16::gfp*) or three (for DAF-2::degron; *daf-16::gfp*) independent experiments. ****: p < 0.0001. (C) Dauer alae of *daf-2(e1370)* and DAF-2::degron animals. Arrows indicate dauer alae specific prongs. Bar = 100 μm. (D) Percentages of animals that were positive for red fluorescent beads in the intestine. n > 100 animals from 3 independent experiments. Error bar represents s.d. ****: p < 0.0001 (E) Images of control and dauer pharynxes. The pharynx of DAF-2::degron animals on auxin were indistinguishable from other *daf-2* mutant dauers. Black arrows indicate wild-type pharynx, and white arrows indicate dauer-typical pharynx constriction. Bar = 50 μm. For (A, D), see Data Source File 1 for raw data and statistics.

**Supplementary Figure 3:**
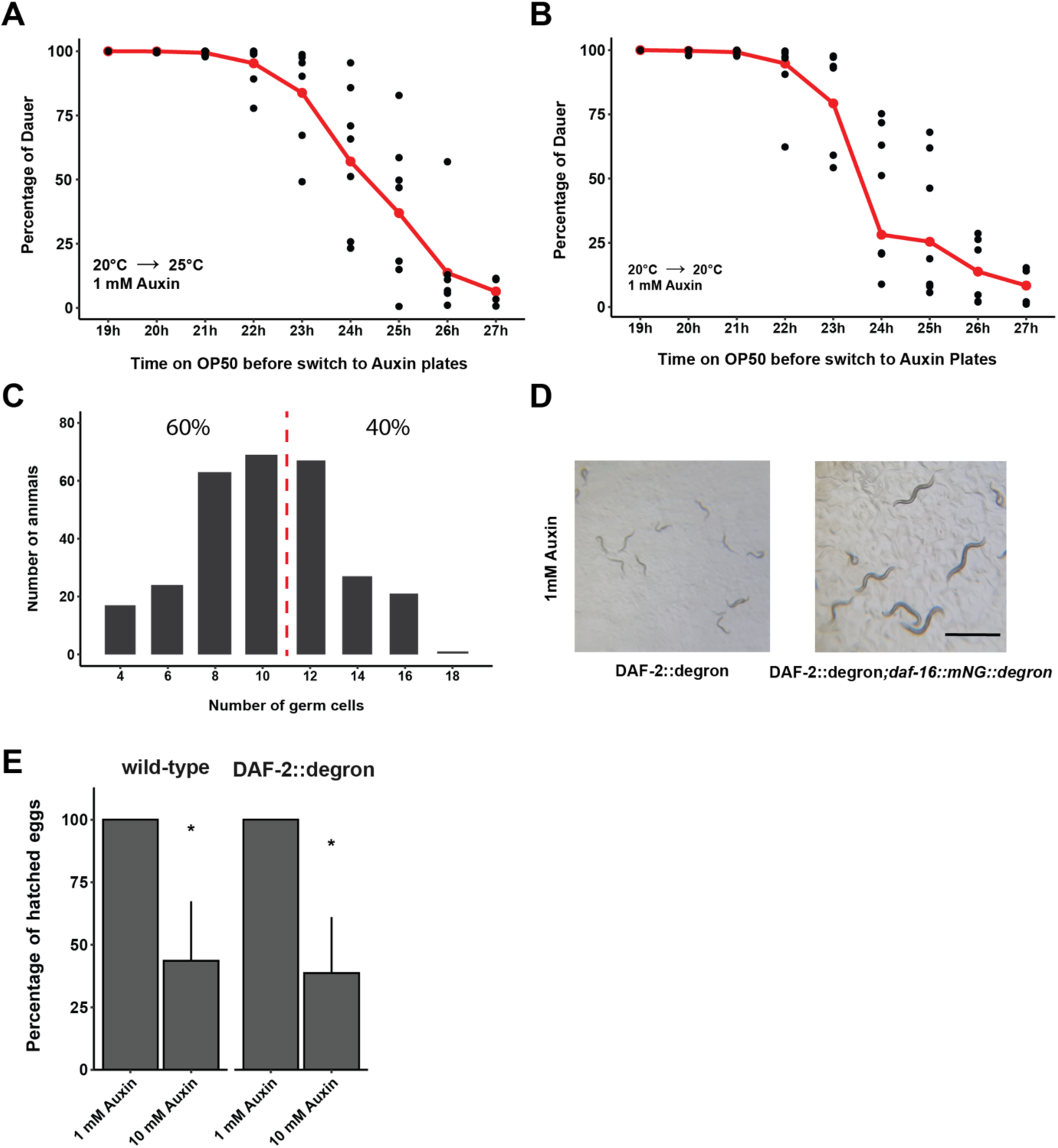
Depletion of DAF-2::degron at mid-L1 promoted *daf-16*-dependent dauer entry. (A) Determination of the decision time point between dauer entry or continuation of normal development of DAF-2::degron animals at 25°C. The animals were synchronized for two days in M9 at 20°C, then put onto plates at 20°C. Then DAF-2::degron animals were put on plates supplemented with Auxin after the indicated time on the x-axis and shifted to 25°C. The ratio of dauers and adult animals was determined two days later. Each time point was repeated 4 to 7 times with n = 976 to 2022. The red line shows the average, black dots indicate single measurements. (B) Determination of the decision time point between dauer entry or continuation of normal development of DAF-2::degron animals at 25°C. The animals were synchronized for two days in M9 at 20°C, then put onto plates at 20°C. Next, DAF-2::degron animals were put on plates supplemented with Auxin after the indicated time on the x-axis and shifted to 20°C. The ratio of dauers and adult animals was determined 3 days later. Each time point was repeated 4 to 7 times with n = 735 to 1791. The red line shows the average, black dots indicate single measurements. Note that dauers at 20°C tend to crawl off the plate, the graph is therefore only an approximation, and there are likely more dauers on the plate. (C) Germ cell count of L1 after 24 to 25 hours on OP50. The dotted red line indicates a younger population that enters dauer and an older population that continues development. The percentages indicate two arbitrarily divided groups with few and many germ cells to match the observation in Figure 3a that approximately 50% of the animals continue development at 24 to 25 hours. The experiment was repeated 5 times with n = 289. (D) Simultaneous knockdown of degron-tagged DAF-16 in DAF-2::degron*; daf-16::mNG::degron* led to fertile adults. Bar = 2 mm. (E) 10 mM Auxin causes embryonic lethality. The same amount of egg suspension was put on plates, and the number of offspring was counted several days later. This experiment was repeated four times for DAF-2::degron and two times for wild type. Error bar represents s.d. *: p < 0.05 For (A-C, E), see Data Source File 1 for raw data and statistics.

**Supplementary Figure 4.**
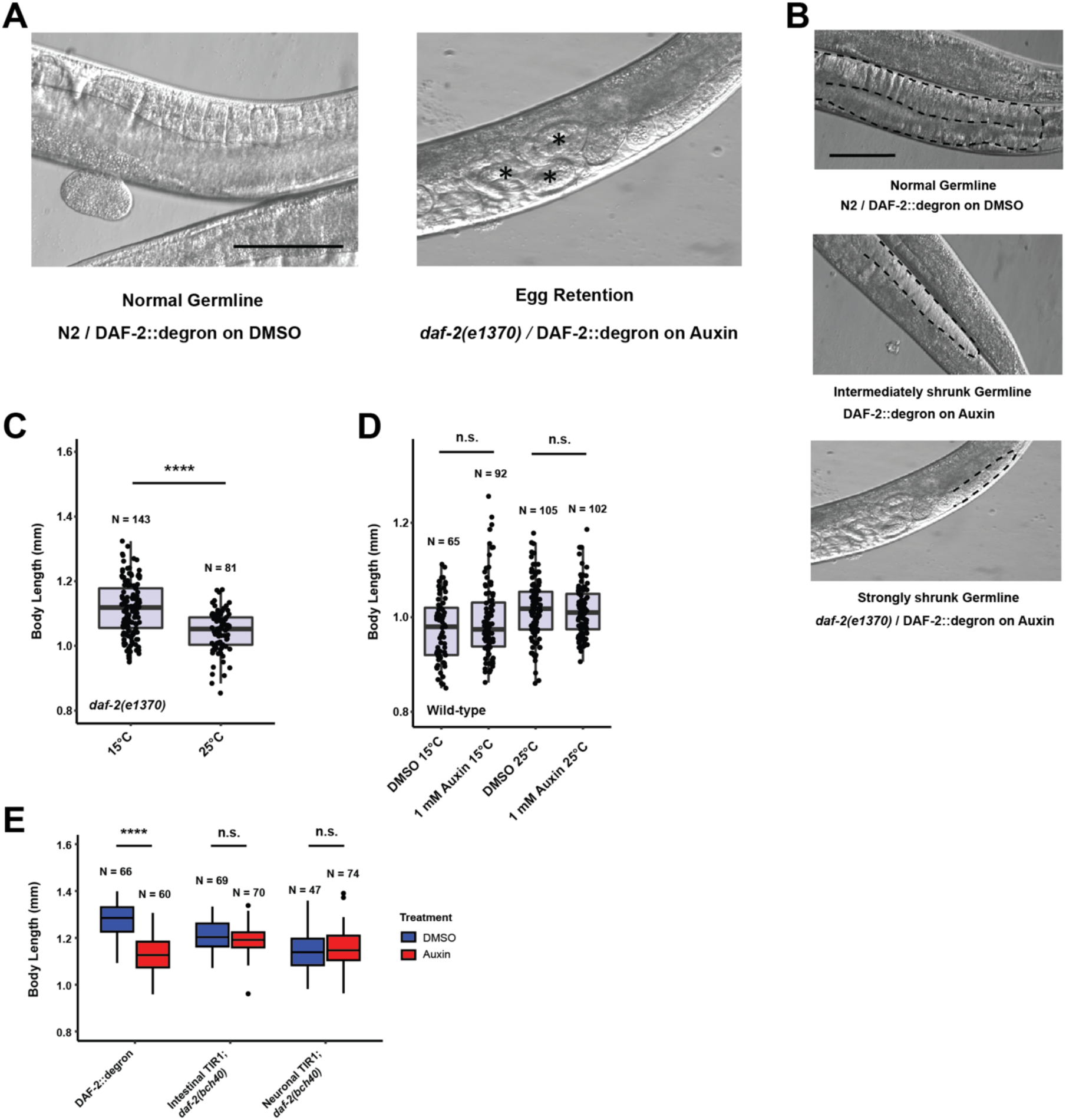
Depletion of DAF-2::degron caused dauer-associated phenotypes at 15°C. (A) Representative images of the egg retention phenotype. The asterisk indicates eggs that are stored in the germline. Bar = 100 μm (B) Representative image of the different levels of germline sizes. Bar = 100 μm (C) *daf-2(e1370)* have reduced body size at 25°C. The experiment was performed 3 independent times. ****: p < 0.0001 (D) Auxin did not affect body size in wild type (N2). The experiment was performed 3 independent times. (E) Depletion of intestinal or neuronal DAF-2::degron did not reduce body size. The experiment was performed two independent times. ****: p < 0.0001 For (C-E), see Data Source File 1 for raw data and statistics.

**Supplementary Figure 5.**
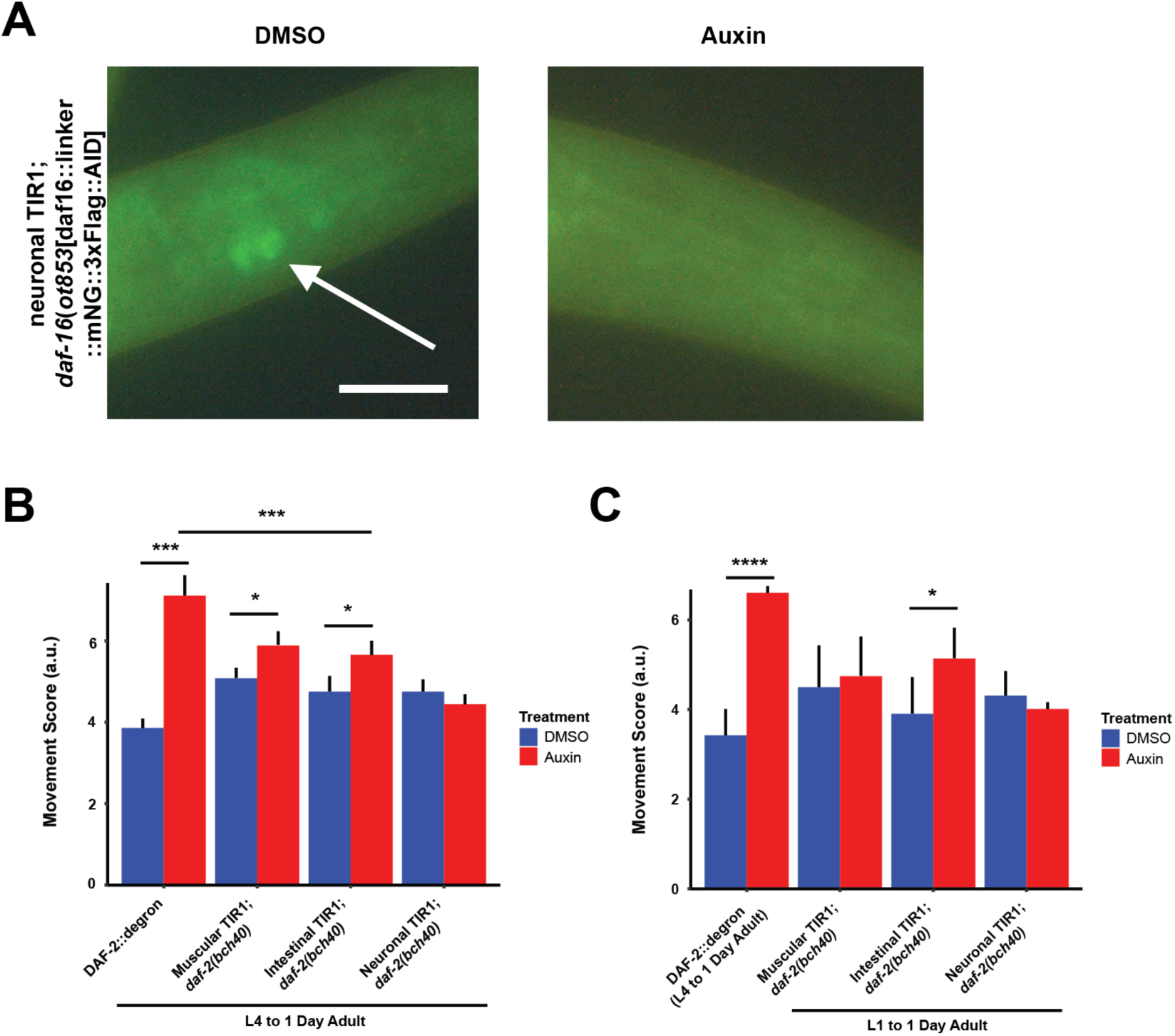
Oxidative resistance and tissue-specific effects after DAF-2 depletion. (A) *daf-16(ot853* [*daf16*::linker::mNG::3xFLAG::AID] was degraded by neuronal TIR1. Arrow indicated neuronally expressed mNeonGreen-and degron-tagged DAF-16. Animals were put for 24 hours on plates with DMSO or 1 mM Auxin, respectively. Bar = 20 μm. (B) Quantification of 1 mM auxin treatment from L4 to first day of adulthood enhanced the arsenite-induced oxidative stress resistance in DAF-2::degron, muscular TIR1; *daf-2(bch40)*, and intestinal TIR1;*daf-2(bch40)*. The experiment was performed 4 independent times (except 3 times for DAF-2::degron). Error bar represents s.d. *: p < 0.05, ***: p < 0.001. (C) Quantification of 1 mM auxin treatment from L1 to the first day of adulthood enhanced the arsenite-induced oxidative stress resistance in muscular TIR1; *daf-2(bch40)* and intestinal TIR1; *daf-2(bch40)*. DAF-2::degron was treated from L4 to the first day of adulthood to bypass dauer entry. The experiment was performed 3 independent times. Error bar represents s.d. *: p < 0.05, ****: p < 0.0001 For (B-C), see Data Source File 1 and 3 for raw data, statistics, and additional trials.

## Materials and Methods

### Strains

All strains were maintained on NGM plates and OP50 *Escherichia coli* at 15°C as described. The strains and primers used in this study can be found in Supplementary Table 3.

### Statistical analysis and plotting

Statistical analysis was either done by using RStudio or Excel. All plots have been made using RStudio (1.2.5001). The packages ggplot2, survminer, dplyr were required for some plots.

### Auxin plates

Auxin (3-Indoleacetic acid, Sigma #I3750) was dissolved in DMSO to prepare a 400 mM stock solution and stored at 4°C. Auxin was added to NGM agar that has cooled down to about 60°C before pouring the plates (Zhang et al., 2015). For lower concentrations (1 μM and 10 μM), the 400 mM stock dilution was further diluted in DMSO. Control plates contained the same amount of DMSO (0.25% for 1 mM auxin plates). For lifespans, plates were supplemented with FUDR for a final concentration of 50 μM.

### Degron-tagged *daf-2* strain and tissue-specific TIR1 expression

The sgRNA targeting the terminal exon of the annotated daf-2 isoform a, 5’ (G)TTTGGGGGTT<u>TCA</u>GACAAG 3’ was cloned into the PU6:: sgRNA (F+E) plasmid backbone, pIK198 (Katic et al., 2015), yielding plasmid pIK323. The initial guanine was added to aid transcription of the sgRNA. The underlined nucleotides in the sgRNA correspond to the stop codon of the DAF-2(A) protein.

The CRISPR tag repair template pIK325 degron::3xFLAG::SL2::SV40::NLS::degron ::wrmscarlet::egl-13 NLS was assembled using the SapTrap method (Schwartz and Jorgensen, 2016) from the following plasmids: pMLS257 (repair template only destination vector), pIK320 (wrmscarlet::syntron-embedded LoxP-flanked, reverse Cbr-unc-119), pMLS285 (egl-13 NLS N-tagging connector), pIK321 (linker::auxin degron::3XFLAG::SL2 operon::SV40 NLS::linker::auxin degron C-tagging connector) and phosphorylated pairs of hybridized oligonucleotides oIK1182 5’TGGTCGGCTTTCGGTGAAAATGAGCATCTAATCGAGGATAATGAGCATCATCCACTTGTC 3’, oIK1183 5’CGCGACAAGTGGATGATGCTCATTATCCTCGATTAGATGCTCATTTTCACCGAAAGCCGA 3’, and oIK1184 5’ACGAACCCCCAAAAAATCCCGCCTCTTAAATTATAAATTATCTCCCACATTATCATATCT 3’, oIK1185 5’TACAGATATGATAATGTGGGAGATAATTTATAATTTAAGAGGCGGGATTTTTTG GGGGTT 3’, respectively. Modules pIK320 and pIK321 (this study), compatible with the SapTrap kit, were assembled through a combination of synthetic DNA (Integrated DNA technologies) and molecular cloning methods. EG4322 *ttTi5605*; *unc-119(ed3)* animals were injected with a mix consisting of the sgRNA pIK323 at 65 ng/ml, tag repair template pIK325 at 50 ng/ml, pIK155 P*eft-3*::Cas9::*tbb-2* 3’UTR at 25 ng/ml and fluorescent markers pIK127 P*eft-3*::GFP::h2b::*tbb-2* 3’UTR at 20 ng/ml and P*myo-3*::GFP at 10 ng/ml. Among the non-Unc F2 progeny of the injected animals not labeled with green fluorescence were correctly tagged *daf-2*(*bch40* [degron::3xFLAG::SL2::SV40 NLS::degron ::wrmscarlet::*egl-13* NLS]) animals. We were able to recover two independent CRISPR alleles *daf-2(bch39)* and *daf-2(bch40)*. pIK280 (TIR1::mRuby::*tbb-2* in a MosSCI-compatible backbone) (Frøkjaer-Jensen et al., 2012; 2008) was created by Gibson assembly (Gibson et al., 2009) from templates including pLZ31 (Zhang et al., 2015). Promoter regions were inserted by Gibson assembly of PCR products into pIK280 to express TIR-1::mRuby in different tissues. Such plasmids were injected into EG4322 *ttTi5605*; *unc-119(ed3)* animals (Frøkjaer-Jensen et al., 2012). The strains are IFM160 *bchSi59* [P*myo-3*::TIR1::mRuby::*tbb-2*] II; *unc-119(ed3)*, IFM161 *bchSi60* [P*vha-6*::TIR1::mRuby::*tbb-2*] II; *unc-119(ed3)*, and IFM164 *bchSi64* [P*rab-3*::TIR1::mRuby::*tbb-2*] II; *unc-119(ed3)*

### Western blot

Synchronized *C. elegans* on their first day of adulthood were shifted to auxin plates for different time points. About 2’000-5’000 Adult *C. elegans* were disrupted using beads in lysis buffer (RIPA buffer (ThermoFisher #89900), 20 mM sodium fluoride (Sigma #67414), 2 mM sodium orthovanadate (Sigma #450243), and protease inhibitor (Roche #04693116001)) and kept on ice for 15 min before being centrifuged for 10 min at 15’000 x g. For equal loading, the protein concentration of the supernatant was determined with BioRad DC protein assay kit II (#5000116) and standard curve with Albumin (Pierce #23210). Samples were boiled at 37°C for 30 min, shortly spun down, and 40 μg of protein was loaded onto NuPAGE Bis-Tris 10% Protein Gels (ThermoFisher #NP0301BOX), and proteins were transferred to nitrocellulose membranes (Sigma #GE10600002). Western blot analysis was performed under standard conditions with antibodies against Tubulin (Sigma #T9026, 1:1000), (Sigma #F3165, 1:1000), FLAG-HRP (Sigma #A8592, 1:1000), and Degron (MBL #M214-3, 1:1000). HRP-conjugated goat anti-mouse (Cell Signaling #7076, 1:2000) secondary antibodies were used to detect the proteins by enhanced chemiluminescence (Bio-Rad #1705061). Quantification of protein levels was determined using ImageJ software and normalized to loading control (Tubulin). Statistical analysis was performed by using either a two-tailed or one-tailed *t*-test. All western blots and quantifications can be found in Data Source File 1 and 2.

### Reporter assays

Transgenic *daf-16::gfp;* DAF-2::degron *C. elegans* were grown on plates for the indicated length of time supplemented with the corresponding concentration auxin at 20°C. For image acquisition, the animals were placed on freshly made 2% agar pads and anesthetized with tetramisole (Teuscher and Ewald, 2018). Images were taken with an upright bright field fluorescence microscope (Tritech Research, model: BX-51-F) and a camera of the model <u>DFK 23UX236</u> (Teuscher and Ewald, 2018). For quantification, the animals were observed under a fluorescent stereomicroscope after the indicated amount of time has passed. *sod-3p::gfp* and *gst-4p::gfp* animals were incubated overnight at 20°C and quantified the next morning. L4 larvae were used for quantification. Statistical analysis was performed using the fisher’s exact test for *daf-16::gfp* and *gst-4::gfp* and two-tailed t-test for *sod-3::gfp*. The DMSO control was compared to the ones treated with various concentrations of auxin.

### Developmental speed

As described in (Ewald et al., 2012), L4 *C. elegans* of wild type N2 and DAF-2::degron were picked to plates at 15°C. After two days, the adult animals were shifted to new plates and were allowed to lay eggs for 2 hours. The stage of the offspring and their health was assayed 4 days later at 20°C. Statistical analysis was performed by using a two-tailed *t*-test.

### SDS dauer assay

Synchronized L1 *C. elegans* were put on 1 mM auxin plates and incubated at 15°C, 20°C, and 25°C, respectively. At the indicated time points, the animals were washed off with M9, shortly centrifuged down, and SDS was added for a final concentration of 1%. After 10 minutes of gentle agitation, the animals were put on plates and checked for survival.

### Dauer pharynx

Adult *C. elegans* were placed on 1 mM auxin plates (for “DAF-2::degron”) or DMSO plates (for “DAF-2::degron”, *daf-2(e1368)* and *daf-2(e1370)*) and shifted to 25°C. Dauer-like offspring or a size-matching control was picked after 4 days, anesthetized in 10 mM sodium azide, and mounted on 2% agarose pads. Images were taken at 40X magnification using an inverted microscope (Tritech Research, MINJ-1000-CUST) and a camera of the model <u>DFK 23UX236</u>.

### Feeding of fluorescent beads

Adult *C. elegans* were put on 1 mM auxin plates, or DMSO plate seeded with OP50 containing a 1:100 dilution of red fluorescent latex bead solution (Sigma #L3280) and shifted to 25°C. Dauer offspring and control L2/L3 on the bacterial lawn were picked after 4 days, anesthetized in 10 mM sodium azide, and mounted on 2% agarose pads. An upright bright field fluorescence microscope (Tritech Research, model: BX-51-F) and a camera of the model DFK 23UX236 were used for image acquisition. The presence of beads in the intestine was checked at 20X magnification. A two-tailed *t*-test was used for analysis.

### Dauer transition assay

Bleached eggs were synchronized for two days at 20°C in M9 buffer supplemented with 5μg/ml cholesterol to yield a highly synchronous population. The larvae were put on OP50 and switched to plates containing 1 mM auxin at different time points. Dauer and non-dauer animals were counted after two days at 25°C or three days at 20°C.

### Germ cell count in L1

Bleached eggs were synchronized for two days at 20°C in M9 buffer supplemented with 5μg/ml cholesterol to yield a highly synchronous population. The larvae were put on OP50 for 24 to 25 hours, washed off and anesthetized with 0.25 mM tetramisole, and mounted on 2% agarose pads. An upright bright field fluorescence microscope (Tritech Research, model: BX-51-F) was used to count the germ cells.

### Germline morphology and egg retention

L4 *C. elegans* maintained at 15°C were picked on 1 mM auxin or control plates and shifted to the indicated temperatures. On the second day of adulthood, animals were mounted on 2% agar pads and anesthetized with 0.25 mM tetramisole. Images were taken at 40X magnification on an inverted microscope (Tritech Research, MINJ-1000-CUST) and a camera of the model <u>DFK 23UX236</u>. Statistical analysis was performed using a two-tailed *t*-test for egg retention and fisher’s exact test for germline morphology.

### Body length measurement

L4 *C. elegans* maintained at 15°C were picked on 1 mM auxin or control plates and shifted to the indicated temperatures. On the second day of adulthood, animals were mounted on 2% agar pads and anesthetized with 0.25 mM tetramisole. Images were taken at 10X magnification with a upright bright field fluorescence microscope (Tritech Research, model: BX-51-F) and a camera of the model DFK 23UX236. Body lengths were measured by placing a line through the middle of the body, starting from head to tail, using ImageJ 1.51j. Statistical analysis was performed by using a two-tailed *t*-test.

### Progeny count

L4 *C. elegans* maintained at 15°C were picked on 1 mM auxin or control plates and shifted to the indicated temperatures. *C. elegans* were shifted when necessary on fresh plates, and the progeny was counted after two days of development. Animals that crawled off the plate, dug into the agar, or bagged precociously were censored. Statistical analysis was performed by using a two-tailed *t*-test.

### Lifespan assays

Synchronized L1 *C. elegans* were cultured at 15°C or 20°C on OP50 and shifted at the L4 stage to NGM plates containing 50 μM FUDR and auxin or DMSO. Bursted, dried out, or escaped animals were censored, and animals were considered dead when they failed to respond to touch and did not show any pharyngeal pumping. For late-life auxin lifespan assays. L4 *C. elegans* were picked onto NGM plates containing 50 μM FUDR. At day 20 or 25 of adulthood, plates were top-coated either DMSO or auxin to reach a final concentration of 0.25% DMSO or 1 mM auxin with 0.25% DMSO. Log-rank was used for statistical analysis. The plots were made by using the R-package survminer or JMP 14.1. All statistics can be found in Supplementary Table 1.

### Arsenite assays

Oxidative stress assay was modified from (Ewald et al., 2017). *C. elegans* of the L1 or L4 stage were shifted on auxin or DMSO plates washed off at the indicated time point, incubated with 5 mM sodium arsenite in U-Shaped 96 well plates, and put into the wMicroTracker (MTK100) for movement scoring. For statistical analysis, the area under the curve was measured, and the mean for each run was calculated. Statistical analysis was performed by using a paired sample *t*-test. All plots can be found in Supplementary File 2

### Author contributions

All authors participated in analyzing and interpreting the data. CYE and RV designed the experiments. IK generated degron/FLAG tag into *daf-2* locus and generated TIR1 tissue-specific strains using CRISPR. TP crossed the DAF-2::degron strain and the transgenes into DAF-2::degron strain. CYE performed late-in-life top-coating auxin lifespans. RV performed all other assays. RV and CYE wrote the manuscript in consultation with the other authors.

### Author Information

The authors have no competing interests to declare. Correspondence should be addressed to C. Y. E.

## Acknowledgment

We thank Cyril Statzer for help with analysis of the lifespan and oxidative stress data, Tea Kohlbrenner for help with crossings and taking the alae pictures, Joy Alcedo and Ben Towbin for critical reading and feedback on the manuscript. Some strains were provided by the CGC, which is funded by NIH Office of Research Infrastructure Programs (P40 OD010440). RC was funded through an EMBO Installation Grant, the TEAM/2016-2/11 programme of the Foundation for Polish Science co-financed by the European Union under the European Regional Development Fund, and the Research Council of Norway grant FRIMEDBIO-286499. CYE was funded by the Swiss National Science Foundation grant PP00P3_163898, and RV by the ETH Research Foundation Grant ETH-30 16-2.

